# Inhibition of *Pseudomonas aeruginosa-*secreted protease IV reduces lung inflammation

**DOI:** 10.64898/2026.07.27.739189

**Authors:** Said M. Daboor, Lauren D. Burton, Trilok Neupane, Rhea Nickerson, Ashley Stueck, Carlie L. Charron, Shannen L. Grandy, Zhong Sun, Zui Wang, Tengfei Zhang, Qingping Luo, Christian Lehmann, Juan Zhou, Janet S. Lee, Jason J. LeBlanc, David N. Langelaan, Xiyang Zhang, Zhenyu Cheng

**Author notes:** Correspondences should be addressed to: Said M. Daboor, Xiyang Zhang, and Zhenyu Chen, **Email:**, and. These authors contribute equally (Said M. Daboor, Lauren D. Burton).

## Abstract

The opportunistic pathogen *Pseudomonas aeruginosa* secretes numerous proteases that disrupt host defenses. Among them, the lysyl endopeptidase PrpL has been implicated in virulence, but its role in inflammatory responses has remained unclear. This study shows that purified PrpL activates the AP-1 transcription factors following instillation into mouse lungs, and drives robust production of IL-1β, IL-6, and TNF, as well as clinical symptoms. A proteolytically inactive mutant of PrpL fails to elicit these responses. Structural analysis of the complex of PrpL and its natural inhibitory propeptide (PrpL_PP_) using X-ray crystallography revealed the inhibitory mechanism of PrpL_PP_, which guided the identification of a pre-existing dipeptide inhibitor (LasBi) that blocked PrpL activity and restricted AP-1-driven cytokine induction and clinical symptoms *in vivo*. This study shows that the alkaline protease AprA degrades PrpL_PP_ and functions redundantly with LasB to liberate mature, active PrpL. Our findings: 1) establish PrpL as a key virulence factor that triggers AP-1-mediated inflammatory signaling, 2) provide structural and functional insights into PrpL inhibition, and 3) identify AprA as a novel upstream PrpL activator. Together, these results highlight PrpL as a promising anti-virulence therapeutic target. Given the functional redundancy of inflammatory effectors produced by *P. aeruginosa*, strategies aimed at mitigating *P. aeruginosa*-induced lung inflammation through PrpL inhibition would likely need to be combined with approaches targeting additional pro-inflammatory bacterial factors.

**Author Summary:** We identify PrpL as a critical *Pseudomonas aeruginosa* protease that activates the inflammatory transcription factor AP-1 and drives potent lung inflammation. Structural analysis of a natural PrpL inhibitor guided development of an existing dipeptide inhibitor that significantly reduced the production of proinflammatory cytokines and clinical responses *in vivo*. Moreover, discovery of the AprA protease as an additional PrpL activator refines the activation model and suggests potential upstream therapeutic strategies. These findings highlight PrpL as both a driver of pathogenesis and a promising therapeutic target for treating *P. aeruginosa* infections.

## Introduction

*Pseudomonas aeruginosa* is a ubiquitous Gram-negative bacterium present in soil and water. It is well-known as an opportunistic pathogen for a wide range of hosts, including plants, nematodes, and vertebrates (1,2). In mammals, *P. aeruginosa* exploits damaged epithelial barriers, impaired mucociliary clearance, and compromised host immune responses to cause infections at various anatomical sites, (e.g., skin, eyes, ears, urinogenital tract, and respiratory system) (1). In people with cystic fibrosis (CF), *P. aeruginosa* causes chronic lung infections that lead to significant morbidity and mortality. *P. aeruginosa* also causes significant acute infections, including ventilator-associated pneumonia and skin and soft tissue infections, underscoring the need to elucidate the underlying mechanisms by which it establishes acute infections (3).

Innate immunity plays a critical role in defense against *P. aeruginosa* infections. In mammals, microbe-associated molecular patterns (MAMPs) from *P. aeruginosa* are first detected by pattern recognition receptors expressed by epithelial and sentinel cells, such as Toll-like receptors and nucleotide-binding oligomerization domain (NOD)-like receptors (4,5). These detection systems activate immune signaling pathways that lead to the assembly and translocation of transcription factors to the cell nucleus, including activator protein 1 (AP-1) and nuclear factor kappa B (NF-κB) (4,6), and also trigger the formation of the inflammasome (5). These transcription factors induce the expression of major inflammatory cytokines, such as interleukin (IL)-6, IL-1β, and tumor necrosis factor (TNF) (6). However, the production of these proinflammatory cytokines by *P. aeruginosa* infection can also lead to excessive lung inflammation and tissue injury (7–9).

Secreted proteases are key contributors to *P. aeruginosa*’s virulence, particularly in acute infections, although they are also present in chronic CF infections (10). One of these virulence factors is protease IV (PrpL), a type 2 secreted serine protease whose expression and secretion are controlled by the Las and Rhl quorum sensing systems (11,12). A recent study showed that transcription of *prpL* is thermoregulated by LasR during stationary phase, but this was not seen during exponential phase (13). PrpL is known to degrade multiple host defense proteins, including plasmin, fibrinogen, immunoglobulin, complement proteins C3 and C1q (14), IL-22 (15), and the surfactant proteins SP-A and SP-D (16). PrpL has been implicated as a major virulence factor in corneal infections, where PrpL-deficient strains exhibited markedly reduced virulence compared to wild-type strains (17). Additionally, PrpL exacerbates infections by other pathogens; for example, co-administration of PrpL with *Streptococcus pneumoniae* in a murine lung infection model led to sepsis and systemic infection (18). These findings highlight PrpL as a critical virulence factor and a potential therapeutic target. However, PrpL has not been shown to directly influence inflammatory signaling pathways in the context of lung infection. Moreover, existing serine protease inhibitors that target PrpL, such as tosyl-L-lysine chloromethyl ketone (TLCK) (14), are highly toxic (19). Critically, there is a lack of safe and effective PrpL inhibitors that can potentially be developed for therapeutic use.

We hypothesized that the proteolytic activity of PrpL directly contributes to the induction of inflammatory signaling pathways in the lung and that inhibition of PrpL could attenuate these responses. To address this hypothesis, we investigated the inflammatory effects of PrpL in mouse lungs and examined the signaling pathways involved. In parallel, we sought to better understand the molecular mechanisms governing PrpL activity by determining the structure of PrpL in complex with its cognate propeptide inhibitor (PrpL_PP_), identifying small-molecule inhibitors of PrpL, and investigating the proteolytic processes involved in PrpL activation. Together, these studies were designed to better define the role of PrpL in host-pathogen interactions, elucidate the mechanisms governing its activity, and provide a foundation for future investigations into strategies for modulating PrpL function.

## Results

### PrpL instillation into the mouse lung elicits strong inflammatory responses

To investigate the effects of PrpL on mammalian inflammatory responses, we instilled purified PrpL, along with its proteolytically inactive mutant form (PrpL^S198A^, which contains a serine 198 to alanine mutation in one of the catalytic triad amino acid residues), intratracheally into mice. LPS was used as a positive control. Both PrpL and PrpL^S198A^ were purified from native *P. aeruginosa* culture supernatant as previously described (20) and were confirmed to contain only negligeable trace amount of endotoxin (1-6 EU/mL and <10pg in any experiments). The purified PrpL was also free of other proteases (i.e. AprA or LasB) (Supplementary Fig. 1).

To establish an appropriate dose of purified PrpL for the intratracheal instillation model, we initially evaluated two doses (0.1 and 0.5 µg/g body weight) for their induction of clinical symptoms and inflammatory responses compared to a commonly used dose (21) of LPS (5 µg/g body weight) (Supplementary Fig. 2). This approach allowed us to determine a PrpL dose that produced a robust yet experimentally manageable inflammatory response. Our results showed that even the lower dose of PrpL (0.1 µg/g body weight) elicited a substantial inflammatory response, supporting the conclusion that PrpL is capable of activating host inflammatory signaling at relatively low concentrations. However, the magnitude and consistency of the response were lower than those observed with the 0.5 µg/g body weight dose (Supplementary Fig. S2). Based on these findings, we selected 0.5 µg PrpL/g body weight for subsequent experiments.

Over the course of all instillations, we examined markers of inflammation, including transcription factor activation, cytokine production, and clinical symptoms. At the transcription factor level, PrpL but not PrpL^S198A^ significantly increased phospho-c-Fos, part of the AP-1 complex (Fig. 1A). Moreover, PrpL increased the phosphorylation of c-Jun, also part of the AP-1 complex, by 5-fold (p = 0.056). Neither form of PrpL increased phospho-IκB levels, whereas LPS induced a significant increase in phosphor-IκB levels 24 hours after instillation (Fig. 1A). Using ELISA, we showed that PrpL but not PrpL^S198A^ instillation elicited a strong induction of three major proinflammatory cytokines (IL-1β, IL-6, and TNF) in both lung (Fig. 1B) and bronchoalveolar lavage fluid (BALF) (Fig. 1C), comparable to the levels observed with LPS instillation. We recorded weight loss at 4 and 24 hours after the course of all instillations, and clinical scores were assigned based on the criteria described in Supplementary Table 1. Both measures served as additional indicators of PrpL’s proinflammatory roles. At their respective doses, LPS and PrpL instillation led to mild symptoms (average clinical scores ∼5) and substantial weight loss of up to ∼15% by 24 hours after instillation (Fig. 1D). By contrast, PrpL^S198A^-instilled mice behaved similarly to PBS control mice for all outcomes measured (Fig. 1D).

**Figure 1.**
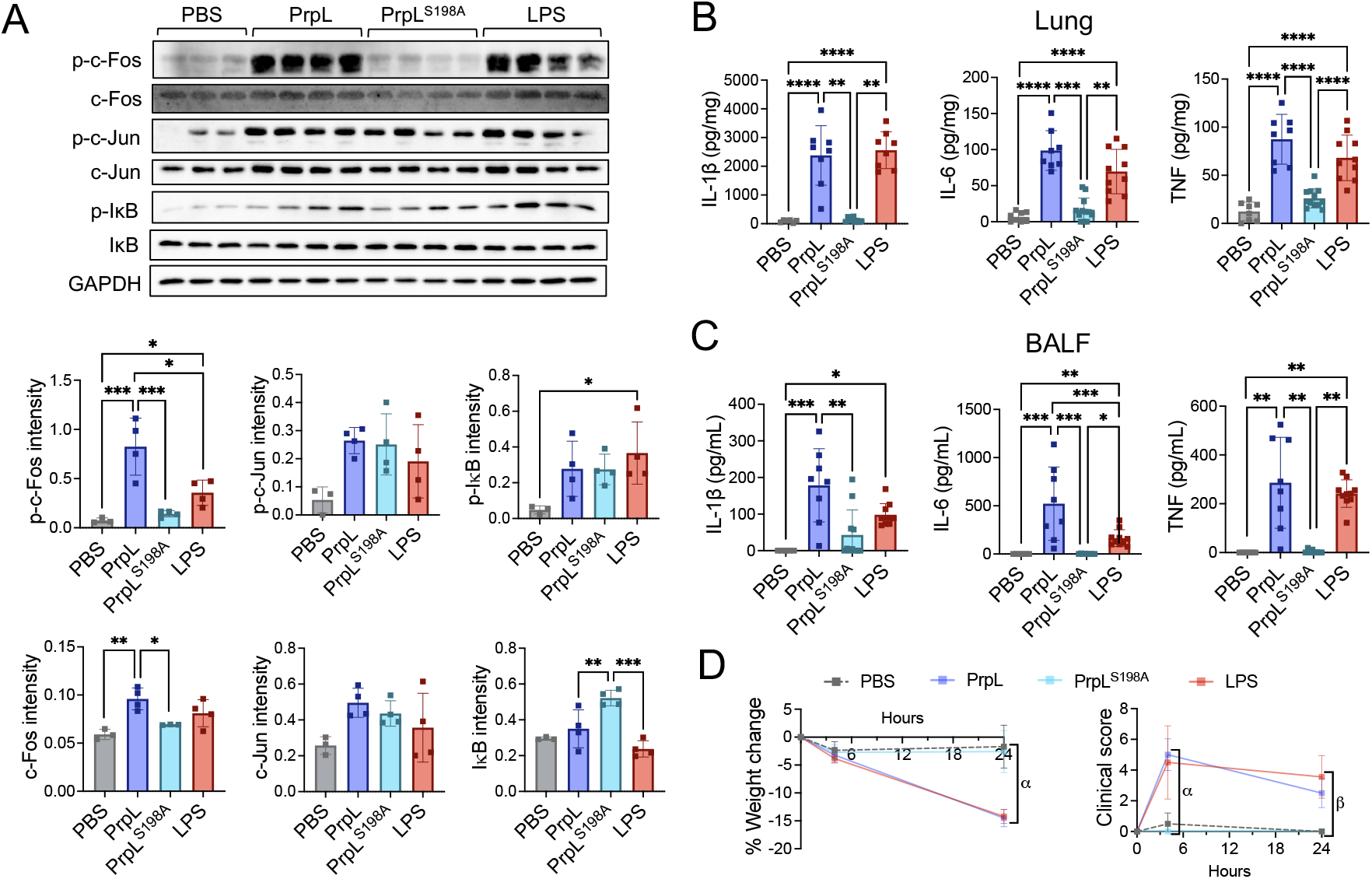
PrpL enzymatic activity is required to activate inflammatory responses *in vivo*. A) Western blots detecting total and phosphorylated (p-) c-Fos, c-Jun, and IκB in lung homogenate. Quantification of band intensity for the blots were normalized to GAPDH loading control. Each lane represents a lung homogenate sample from an individual mouse, with different lanes corresponding to different mice. B) Concentrations of inflammatory cytokines IL-1β, IL-6, and TNF in lung homogenate, expressed per mg total lung protein. C) Concentrations of IL-1β, IL-6, and TNF in BALF, expressed per mL BALF. D) Percentage change in mouse body weight measured at 4- and 24-hours post-instillation of PrpL or LPS, relative to body weight pre-instillation. α represents the following comparisons: PrpL vs. PBS = ****, LPS vs. PBS = ****, PrpL vs. PrpL^S198A^ = ****. Clinical scores assigned at 4- and 24-hours post-instillation. α represents the following comparisons: PrpL vs. PBS = ****, LPS vs. PBS = ****, PrpL vs. PrpL^S198A^ = ****; and β represents the following: PrpL vs. PBS = ****, LPS vs. PBS = ****, PrpL vs. PrpL^S198A^ = ****, PrpL vs. LPS = *. Data are represented as mean ± standard deviation and were compared by one-way ANOVA with correction for multiple comparisons or Mann-Whitney test where applicable. Non-significant = p >0.05, * = p <0.05, ** = p <0.01, *** = p <0.001, and **** = p <0.0001. Comparisons not otherwise indicated are non-significant. *n* ≥10 mice per group, except *n* ≥6 for PBS controls.

We next investigated the proinflammatory impact of PrpL in the context of infection, despite the expectation that redundancies between various bacterial immune modulators might mask some of the effects observed above using purified PrpL instillation. Using an acute lung infection model, we compared host inflammatory responses between the lungs of mice infected with wild-type *P. aeruginosa* (WT PA14) and those infected with a Δ*prpL* deletion mutant. In our model, *P. aeruginosa* infections (both WT and the Δ*prpL* mutant) comparably increased phospho-c-Fos levels (Fig. 2A). It is worth noting that the levels of phospho-c-Fos activation varied among individual infected mice. However, this variability was comparable between mice infected with the WT and Δ*prpL* strains, as both groups exhibited a similar range and magnitude of phospho-c-Fos induction (Fig. 2A). We also did not observe major differences in proinflammatory cytokine production (Fig. 2B & C), clinical scores, or weight loss between PA14 WT and Δ*prpL* deletion mutant-infected mice (Fig. 2D). In addition, there were no significant differences in histological lung inflammatory score or area (Supplementary Fig. 3A-C) or immune cell recruitment (Supplementary Fig. 4A & B) between the WT PA14 and Δ*prpL* deletion mutant-infected mice. Finally, strain-to-strain variation among *P. aeruginosa* isolates can substantially influence infection phenotypes. To address this concern, we generated a Δ*prpL* deletion mutant in the PAO1 background and evaluated it in our *in vivo* infection model. Consistent with our observations using the PA14 strains, the PAO1/Δ*prpL* mutant did not exhibit significant differences in the inflammatory parameters described above compared with its corresponding wild-type strain (Supplementary Fig. 5).

**Figure 2.**
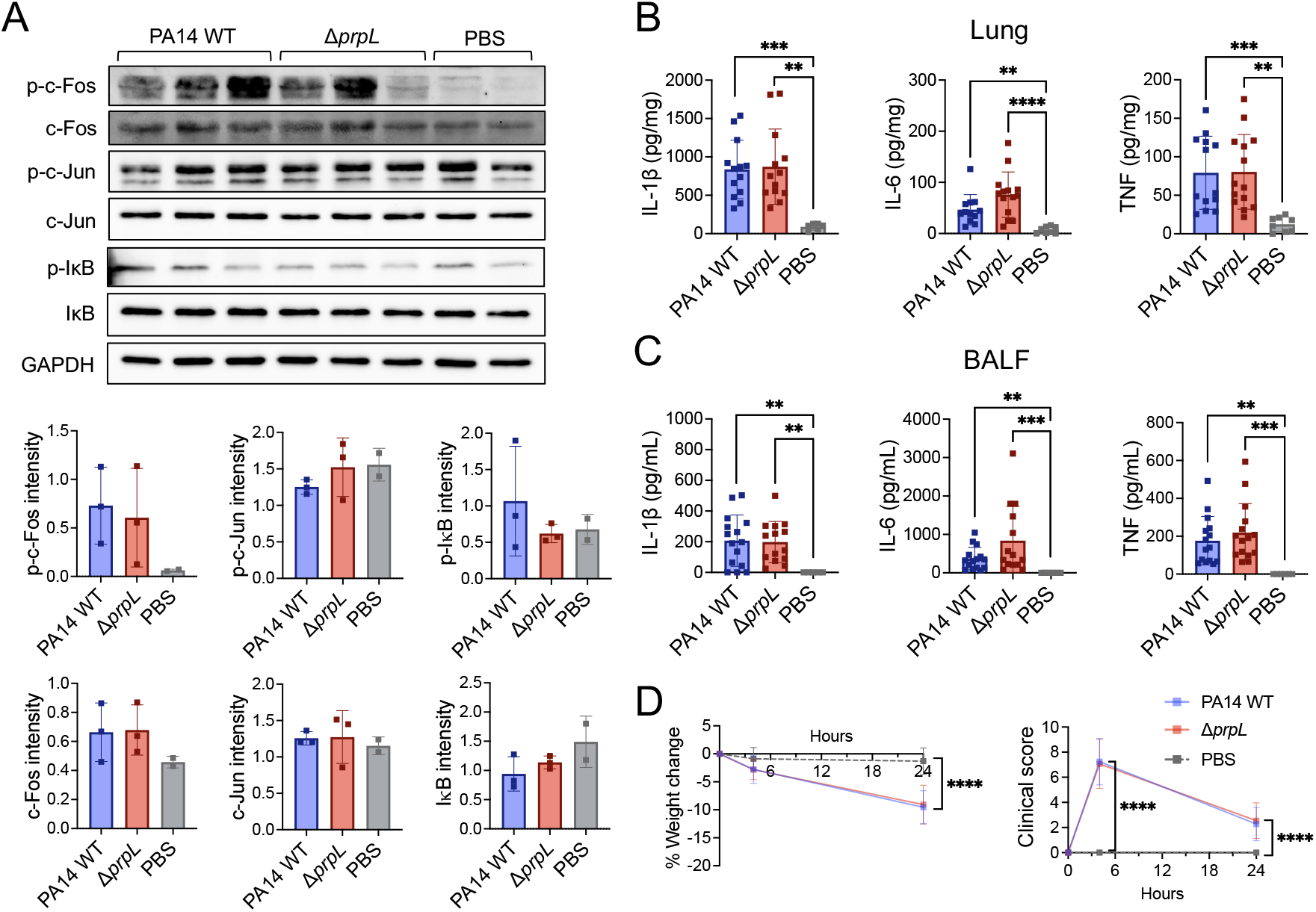
Comparisons of inflammatory responses in mice infected with *P. aeruginosa* PA14 WT or Δ*prpL* mutant. Mice were intratracheally infected with 5×10^5^ CFU of *P. aeruginosa* PA14 wild-type (WT) or Δ*prpL*. Lungs and BALF were collected 24-hours post-infection. A) Western blots detecting total and phosphorylated (p-) c-Fos, c-Jun, as well as IκB in lung homogenate. Quantification of band intensity for the blots were normalized to GAPDH loading control. Each lane represents a lung homogenate sample from an individual mouse, with different lanes corresponding to different mice. B) Concentrations of inflammatory cytokines IL-1β, IL-6, and TNF in lung homogenate, expressed per mg total lung protein. C) Concentrations of IL-1β, IL-6, and TNF in BALF, expressed per mL BALF. D) Percentage change in mouse body weight measured at 4- and 24-hours post-infection. Clinical scores assigned at 4- and 24-hours post-infection. Data are represented as mean ± standard deviation and were compared by one-way ANOVA with correction for multiple comparisons or Mann-Whitney test where applicable. Non-significant, p >0.05, ** = p <0.01, *** = p <0.001, and **** = p <0.0001. Comparisons not otherwise indicated are non-significant. *n* ≥10 per group, except *n* ≥6 for PBS controls.

### X-ray crystallographic analysis of the PrpL-PrpL_PP_ complex reveals the mechanism of PrpL inhibition

Given that PrpL induces AP-1 activation in the host to promote inflammation, we examined the structure of PrpL with the goal of targeting this key virulence factor. PrpL contains an N-terminal signal peptide (SP), a central propeptide, and a C-terminal mature protease (14). The full-length protein has a molecular weight of approximately 48 kDa, while the mature protease and the released propeptide (PrpLpp) are approximately 26 kDa and 19 kDa, respectively (Fig. 3A). The native propeptide inhibitor PrpL_PP_ has been shown to inhibit PrpL (22). There is considerable structural information available for mature lysyl-class proteases, to which PrpL belongs (23). However, structural data on their propeptides, especially in the context of a co-structure, which is crucial for understanding the inhibitory mechanism, has not been reported. To solve the co-structure of the PrpL-PrpL_PP_ complex, we purified native PrpL from *P. aeruginosa* culture media as previously described (20), and recombinant PrpL_PP_ from *E. coli* (Fig. 3B & C). Although PrpL_PP_ inhibits PrpL activity (22), prolonged incubation led to degradation of the PrpL_PP_ protein (Fig. 3D). To address this, we modified our crystallographic approach by using the inactive catalytic triad mutant form, PrpL^S198A^, along with the serine protease inhibitor TLCK. The high-affinity interaction between PrpL and PrpL_PP_ was confirmed using a pull-down assay (Fig. 4A), and the PrpL^S198A^-PrpL_PP_ complex was isolated by size exclusion chromatography (Fig. 4B). SDS-PAGE analysis indicated there was some self-cleavage of PrpL^S198A^ and PrpL_PP_ (Fig. 4A). However, the tertiary structure of the complex was intact as light scattering determined the molecular weight of purified PrpL-PrpL_PP_ to be 45.7 kDa, which is consistent with its expected size (47.8 kDa) (Fig. 4C).

**Figure 3.**
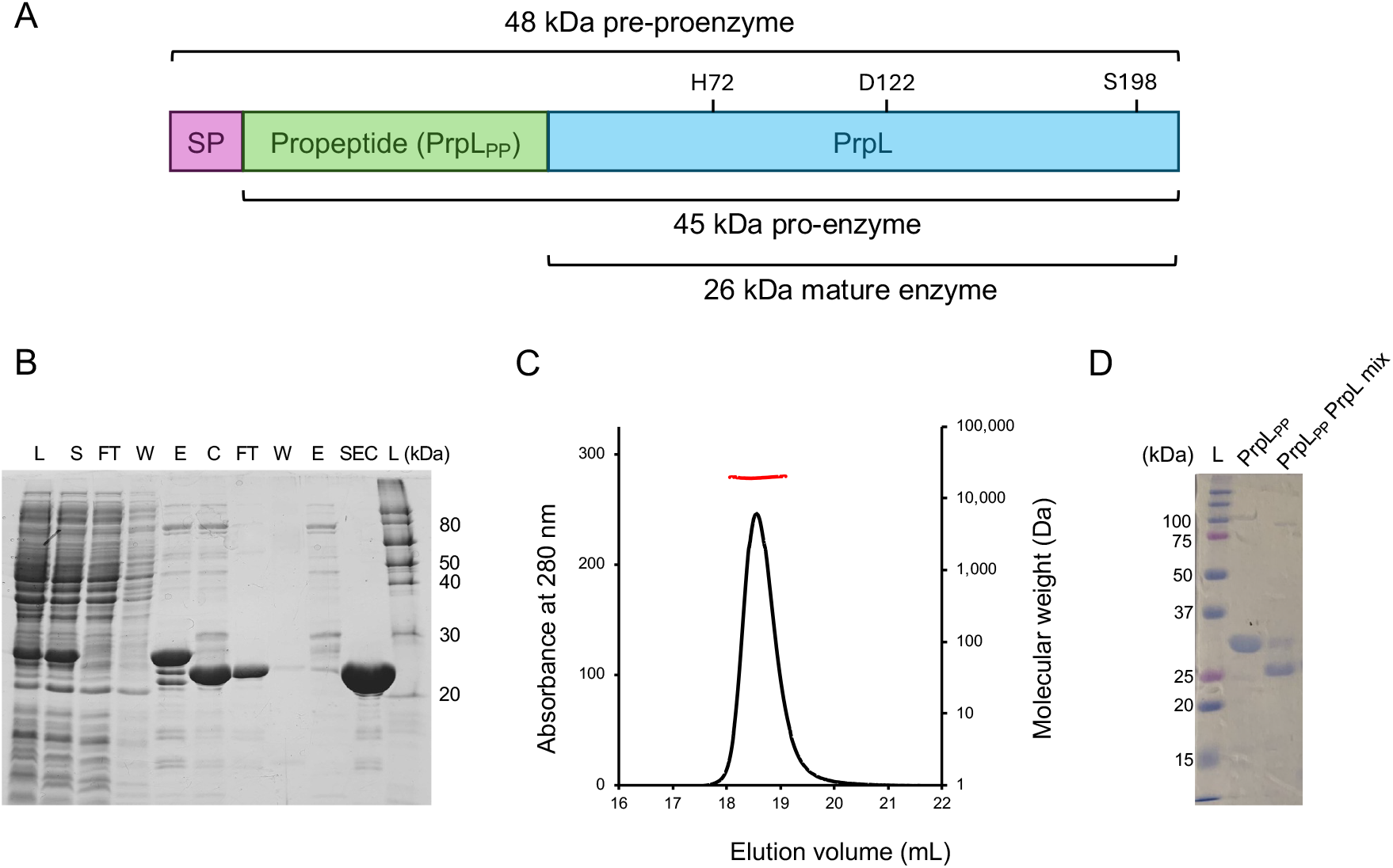
Expression and purification of propeptide (PrpL_PP_). A) Domain structure of PrpL with an N-terminal signal peptide (SP), a central propeptide, and a C-terminal mature protease. The mature PrpL has a catalytic triad consisting of His72, Asp122, and S198 that is required for its catalytic activity. B) Recombinant PrpL_PP_ was expressed in *E. coli* BL21 (DE3) cells, purified by nickel affinity and size-exclusion chromatography (SEC), then analyzed by SDS-PAGE. Gel lanes are L: lysate; S: lysis supernatant; FT: flow-through; W: wash; E: elution; C: after His tag cleavage by TEV protease, SEC: after size exclusion chromatography and L: molecular weight marker. C) Purified propeptide was analyzed by SEC coupled with multi-angle light scattering (MALS). The measured molecular weight of ∼19.4 kDa indicates that isolated PrpL_PP_ exists as a monomer. D) Recombinant PrpL_PP_ (before His tag cleavage) was degraded by PrpL after extended incubation.

**Figure 4.**
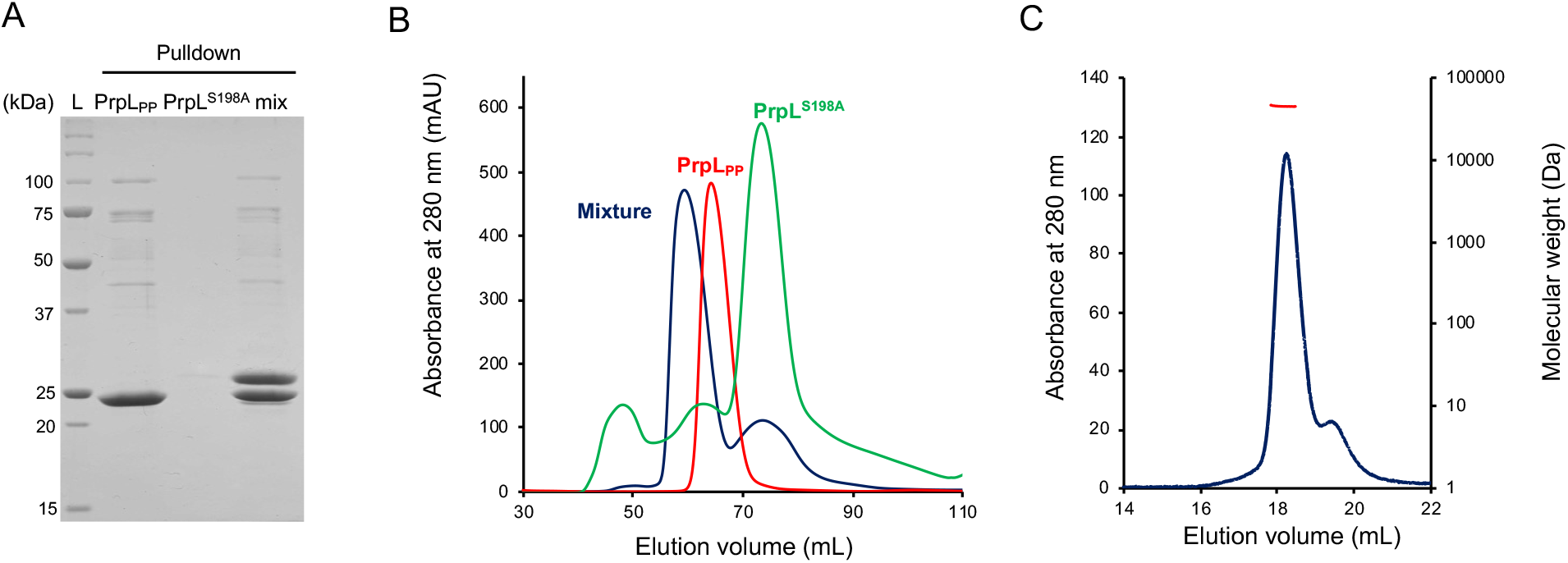
Binding between PrpL and propeptide. A) Binding between propeptide and protease IV was confirmed by incubating Ni^2+^ affinity resin with His_6_-tagged PrpL_PP_, PrpL, or a mixture of both proteins. The presence of both proteins in the mixed sampled confirms binding between PrpL_PP_ and PrpL. B) Size exclusion analysis of PrpL^S198A^, PrpL_PP_, and a mixture of both proteins. The mixed sample elutes earlier than the isolated proteins, indicating the formation of a complex between PrpL_PP_ and PrpL^S198A^. C) Purified PrpL^S198A^-PrpL_PP_ complex was analyzed by SEC-MALS to have a molecular weight of ∼45.7 kDa.

Using a microbatch crystallization setup, crystal formation was observed after several days. Diffraction data was collected and the crystal structure of PrpL^S198A^-PrpL_PP_ was determined to a resolution of 3.3 Å (Table 1). PrpL^S198A^ adopts a trypsin-like fold consisting of two β-barrels that form two closely associated domains, with the catalytic triad located in the cleft between these domains (24,25). PrpL^S198A^ contains three disulfide bridges that are conserved amongst the chymotrypsin family of serine proteases (26). PrpL_PP_ is composed of 8 anti-parallel β-strands divided into two anti-parallel β-sheets that are associated as a jelly-roll beta-sandwich (Fig. 5). The N-terminal 60 amino acids do not adopt a canonical secondary structure. PrpL_PP_ closely associates with PrpL^S198A^ through polar interactions involving β-sheets 2, 4, and 8, while Met31-Ser41 also extends from PrpL_PP_ to form an antiparallel β-sheet with Ala34-Ala43 of PrpL^S198A^. Finally, Lys81-Gly89 extend from PrpL_PP_, form a β-strand with PrpL^S198A^, and occlude its active site. This mechanism of protease inhibition is similar to the inhibition of porcine pancreatic trypsin by the Kunitz-type soybean trypsin inhibitor (27).

**Figure 5.**
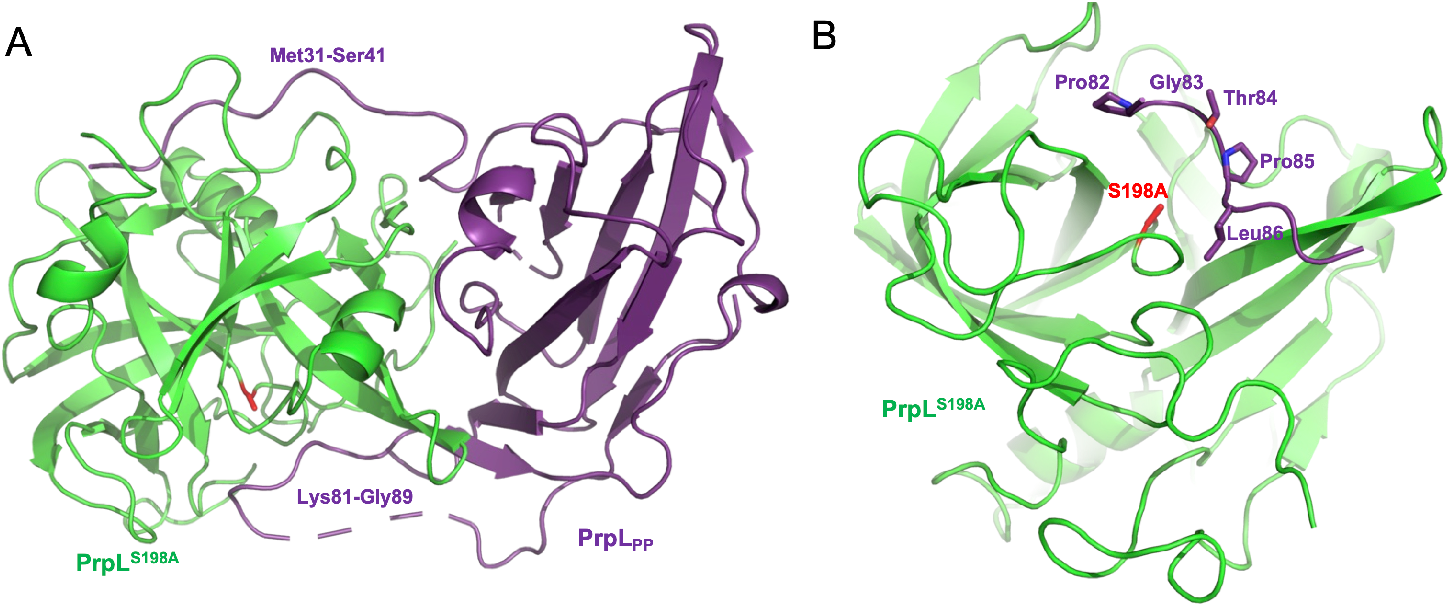
Crystal structure of the PrpL^S198A^-PrpL_PP_ complex. A) Ribbon representation of PrpL^S198A^ (green) bound to PrpL_PP_ (purple). The S198A mutation is shown in red and regions of PrpL_PP_ that form an extended binding interface with PrpL^S198A^ are indicated. B) The active site of PrpL with the S198A mutation is shown in red. Pro82-Leu86 of PrpL_PP_ occlude the active site of PrpL^S198A^ and are shown as sticks.

**Table 1.** Data collection and refinement statistics of the PrpL^S198A^-PrpL_PP_ co-structure.

|  | PrpL<sup>S198A</sup>-PrpL<sub>PP</sub> |
| --- | --- |
| **Wavelength (Å)** | 0.953 |
| **Resolution range** | 48.05 – 3.3 (3.418 – 3.3)<sup>a</sup> |
| **Space group** | P 41 2 2 |
| **Unit cell** | 192.12 192.12 231.84 90 90 90 |
| **Total reflections** | 1766842 (183528) |
| Unique reflections | 65700 (6463) |
| Multiplicity | 26.9 (28.4) |
| Completeness (%) | 99.87 (99.97) |
| Mean I/sigma(I) | 19.02 (3.63) |
| Wilson B-factor | 89.30 |
| R-merge | 0.1914 (1.328) |
| R-meas | 0.1951 (1.352) |
| R-pim | 0.03748 (0.2528) |
| CC1/2 | 0.999 (0.898) |
| CC* | 1 (0.973) |
| Reflections used in refinement | 65657 (6463) |
| Reflections used for R-free | 1282 (127) |
| R-work | 0.2621 (0.3255) |
| R-free | 0.2908 (0.3379) |
| CC(work) | 0.918 (0.758) |
| CC(free) | 0.850 (0.655) |
| Number of non-hydrogen atoms | 23795 |
| macromolecules | 23795 |
| ligands | 0 |
| solvent | 0 |
| Protein residues | 3214 |
| RMS(bonds) | 0.002 |
| RMS(angles) | 0.48 |
| Ramachandran favored (%) | 92.56 |
| Ramachandran allowed (%) | 6.64 |
| Ramachandran outliers (%) | 0.80 |
| Rotamer outliers (%) | 0.00 |
| Clashscore | 7.99 |
| Average B-factor | 89.28 |
| macromolecules | 89.28 |

<sup>a</sup>Statistics for the highest-resolution shell are shown in parentheses.

### Small peptide LasBi inhibits PrpL activity and alleviates inflammation *in vivo*

Based on the PrpL structure, we synthesized the octamer from PrpL_PP_ that is in proximity to the catalytic triad of PrpL. However, the synthetic octamer at 100 µM (maximal solubility) slightly enhanced PrpL activity by ∼16% instead of inhibiting it (Supplementary Fig. 6A). As an alternative strategy, we searched for known stable peptide inhibitors that have shown robust inhibition of other *P. aeruginosa* proteases and may also inhibit PrpL. Hence, we selected the synthetic dipeptide-based inhibitor N-mercaptoacetyl-Phe-Tyr-amide (LasBi), which targets LasB from *P. aeruginosa* (28), for further characterization. With the synthetic dipeptide LasBi, we found a dose-dependent inhibition of PrpL activity, with 46% inhibition at a concentration of 200 μM (Supplementary Fig. 6B). We then incubated PrpL with 200 µM LasBi for 1 hour at 37 °C prior to instillation into mouse lungs. In accordance with the *in vitro* inhibition observed, we found that mice instilled with LasBi-treated PrpL had significantly lowered phospho-c-Fos levels compared to mice treated with PrpL alone. In addition, only PrpL, but not LasBi-treated PrpL, significantly increased phospho-c-Jun levels (Fig. 6A) Importantly, LasBi almost completely abrogated the strong induction of proinflammatory cytokine IL-1β, and significantly lowered both IL-6 and TNF induction by PrpL (Fig. 6B & C). Finally, LasBi-treated PrpL also led to significantly lowered clinical scores at 24 hours post instillation (Fig. 6D).

**Figure 6.**
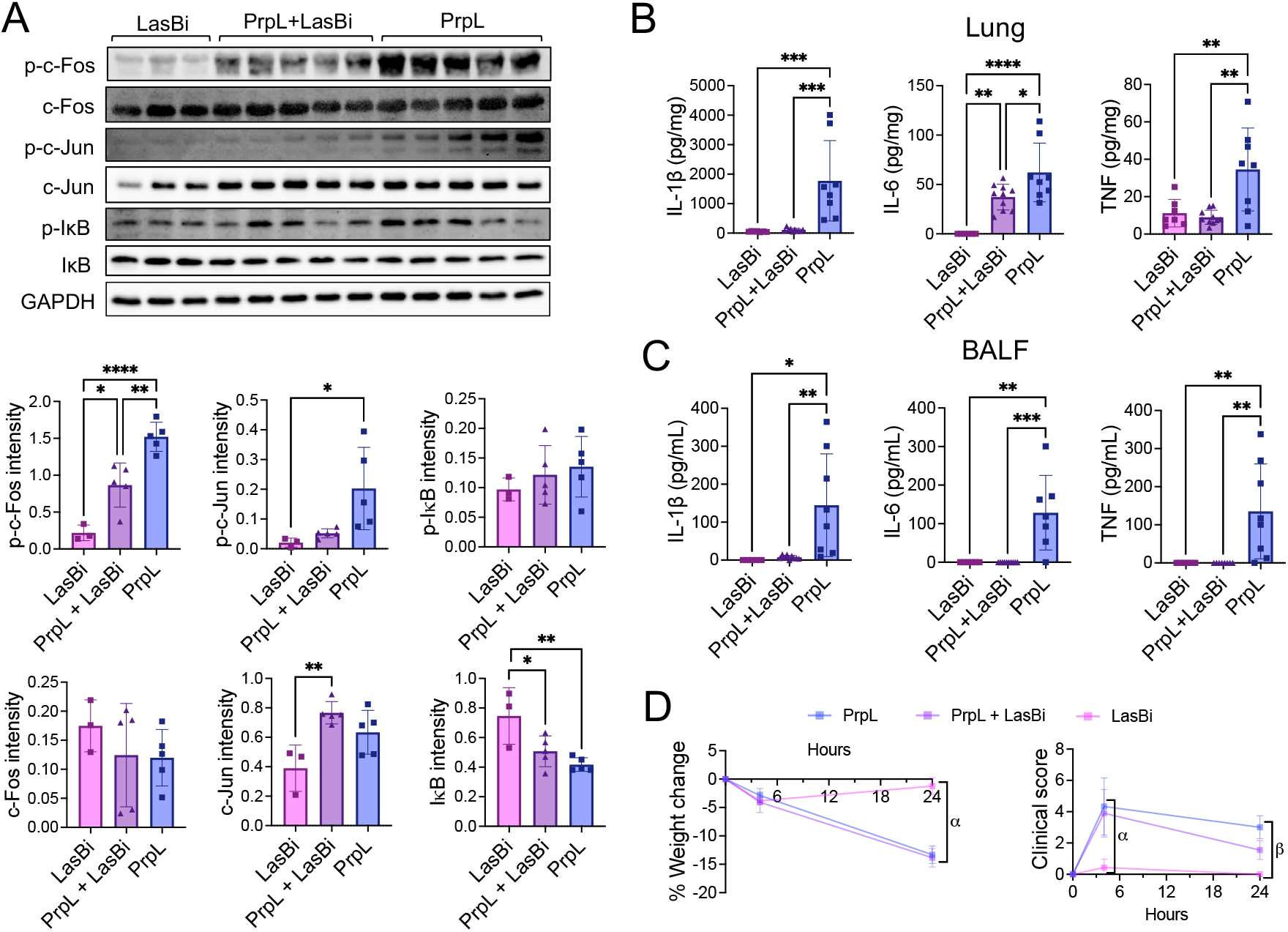
Dipeptide LasBi inhibits PrpL-elicited inflammatory responses *in vivo*. PrpL was incubated with 200 µM LasBi inhibitor for 1 hour at 37°C before being instilled intratracheally into the lungs of mice at a dose of 0.5 µg PrpL per g body weight. Untreated PrpL and LasBi alone served as controls. A) Western blots detecting total and phosphorylated (p-) c-Fos, c-Jun, and IκB in lung homogenate. Quantification of band intensity for the blots were normalized to GAPDH loading control. Each lane represents a lung homogenate sample from an individual mouse, with different lanes corresponding to different mice. B) Concentrations of inflammatory cytokines IL-1β, IL-6, and TNF in lung homogenate, expressed per mg total lung protein. C) Concentrations of IL-1β, IL-6, and TNF in BALF, expressed per mL BALF. E) Percentage change in mouse body weight measured at 4- and 24-hours post-instillation of PrpL and/or LasBi, relative to body weight pre-instillation. α represents the following comparisons: PrpL+LasBi vs. LasBi = ****, LasBi vs PrpL = ****. F) Clinical scores assigned at 4- and 24-hours post-instillation. α represents the following comparisons: PrpL+LasBi vs. LasBi = ****, LasBi vs PrpL = ****; and β represents the following: PrpL vs. PrpL+LasBi = ****, PrpL+LasBi vs. LasBi = ****, LasBi vs PrpL = ****. Data are represented as mean ± standard deviation and were compared by one-way ANOVA with correction for multiple comparisons or Mann-Whitney test where applicable. Non-significant = p >0.05, * = p <0.05, ** = p <0.01, *** = p <0.001, and **** = p <0.0001. Comparisons not otherwise indicated are non-significant. *N* ≥10 per group, except *n* ≥6 for LasBi controls.

### AprA is a novel and potent degrader of PrpL_PP_ and is redundant with LasB in activation of PrpL

In search of other strategies for targeting PrpL activity and virulence, we explored the native maturation and activation process of PrpL in *P. aeruginosa*. It was previously shown that LasB degrades PrpL_PP_ to release active PrpL (22). To further investigate PrpL activation, we examined the supernatants of the WT PA14 strain and several deletion and transposon mutants in the PA14 background for PrpL activity. Surprisingly, the Δ*lasB* mutant supernatant showed the same level of PrpL activity as WT PA14, whereas a Δ*prpL* mutant, a Δ*lasR*-Δ*rhlR* quorum sensing mutant, and a type 2 secretion apparatus *xcpR* transposon mutant all exhibited negligible background levels of activity (Fig. 7A). These data are consistent with another study showing that a LasB-negative *P. aeruginosa* strain PA103 and derivative strains produced considerable quantities of active PrpL (23). This suggests that other proteases may have similar and redundant activities to LasB in degrading PrpL_PP_ to activate PrpL. Previous studies have shown that LasB and the type 1 secreted alkaline protease AprA cooperatively degrade flagellin to suppress MAMP responses in both mammalian cells and plants (29,30). This redundancy likely serves as a safety-net strategy, allowing *P. aeruginosa* to evade host immunity under diverse conditions, as LasB and AprA have different expression patterns and optimal enzymatic conditions (29,30). Indeed, while the supernatant of an Δ*aprA* mutant showed WT-levels of PrpL activity, the Δ*aprA*-Δ*lasB* double mutant showed significantly reduced PrpL activity (Fig. 7A), suggesting redundant functions for AprA and LasB in degrading PrpL_PP_. To detect various forms of PrpL, we generated rabbit polyclonal antibodies against both recombinantly expressed mature PrpL and PrpL_PP_. Consistent with the PrpL activity data, we found that PrpL was fully processed to its mature form in the supernatants of WT and all tested mutants, except in the Δ*aprA*-Δ*lasB* double mutant, where the major species were the 45 kDa PrpL proenzyme (pro-PrpL) and a 37 kDa intermediate, with only trace amounts of the mature form (Fig. 7B). This finding was confirmed using the PrpL_PP_ antibody, which showed no visible PrpL_PP_ within pro-PrpL in the supernatant except in the Δ*aprA*-Δ*lasB* double mutant (Fig. 7C).

**Figure 7.**
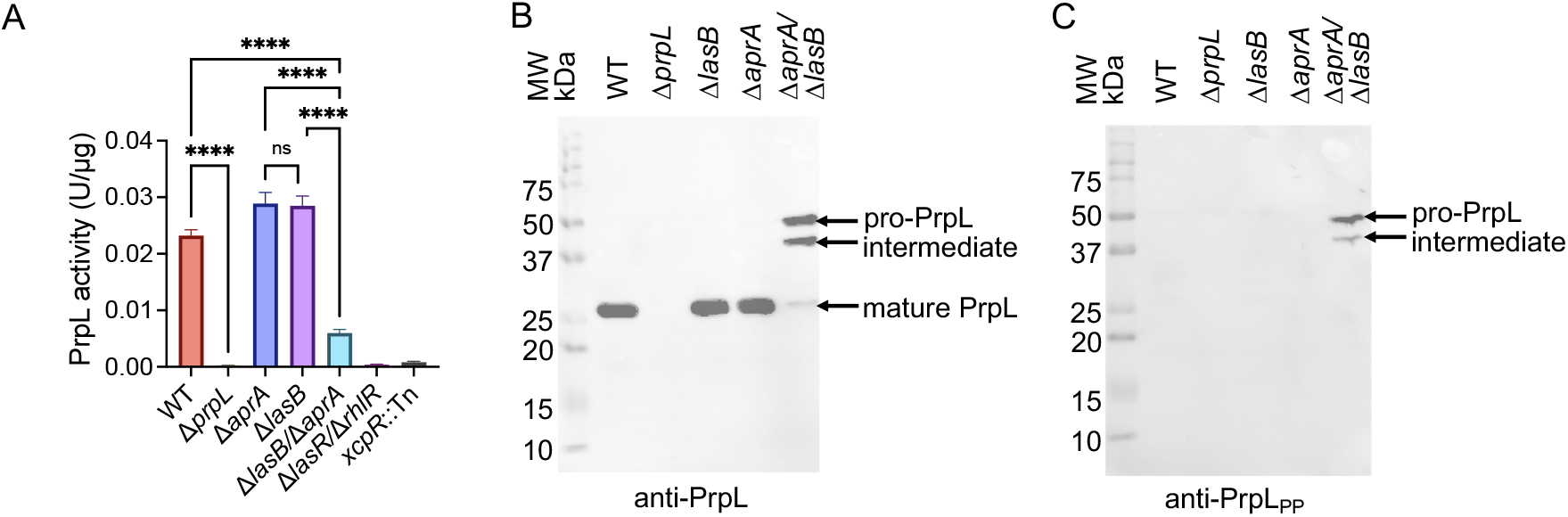
AprA and LasB cooperatively degrade the PrpL propeptide (PrpL_PP_) to activate mature PrpL. PrpL activity assays (A) and western blotting using anti-PrpL (B) and anti-PrpL_PP_ (C) antibodies were performed on equal amounts of protein from culture supernatants of PA14 WT and mutants. The Δ*aprA*Δ*lasB* double mutant exhibited markedly reduced PrpL activity, whereas Δ*aprA* and Δ*lasB* single mutants retained WT-level activity. In all strains except the Δ*prpL* negative control and the Δ*aprA*Δ*lasB* double mutant, PrpL was present predominantly in its mature form, with no detectable free PrpL_PP_. In the Δ*aprA*Δ*lasB* double mutant, PrpL remained largely unprocessed, appearing as pro-PrpL or an intermediate form. Data are represented as mean ± standard deviation and were compared by one-way ANOVA with correction for multiple comparisons. Non-significant = p >0.05 and **** = p <0.0001. Comparisons not otherwise indicated are non-significant.

We then sought to complement the genetic data with biochemical evidence to support this previously unrecognized role of AprA in PrpL_PP_ degradation. We purified native AprA from *P. aeruginosa* supernatant fluid using Fast Protein Liquid Chromatography (FPLC). When purified AprA was mixed with PrpL_PP_ at a ratio of 1:100 and incubated for less than 10 minutes, it led to the complete disappearance of PrpL_PP_ (Fig. 8A). To confirm that the degradation of PrpL_PP_ by AprA was due to specific enzymatic activity, we used the metalloprotease inhibitor phosphoramidon (31) to block AprA activity before mixing it with PrpL_PP_. This inhibitor reduced PrpL_PP_ degradation until up to 5 minutes after incubation. However, even in the presence of phosphoramidon, PrpL_PP_ was still completely degraded after 10 minutes (Fig. 8A), indicating again the high potency of AprA in degrading PrpL_PP_.

**Figure 8.**
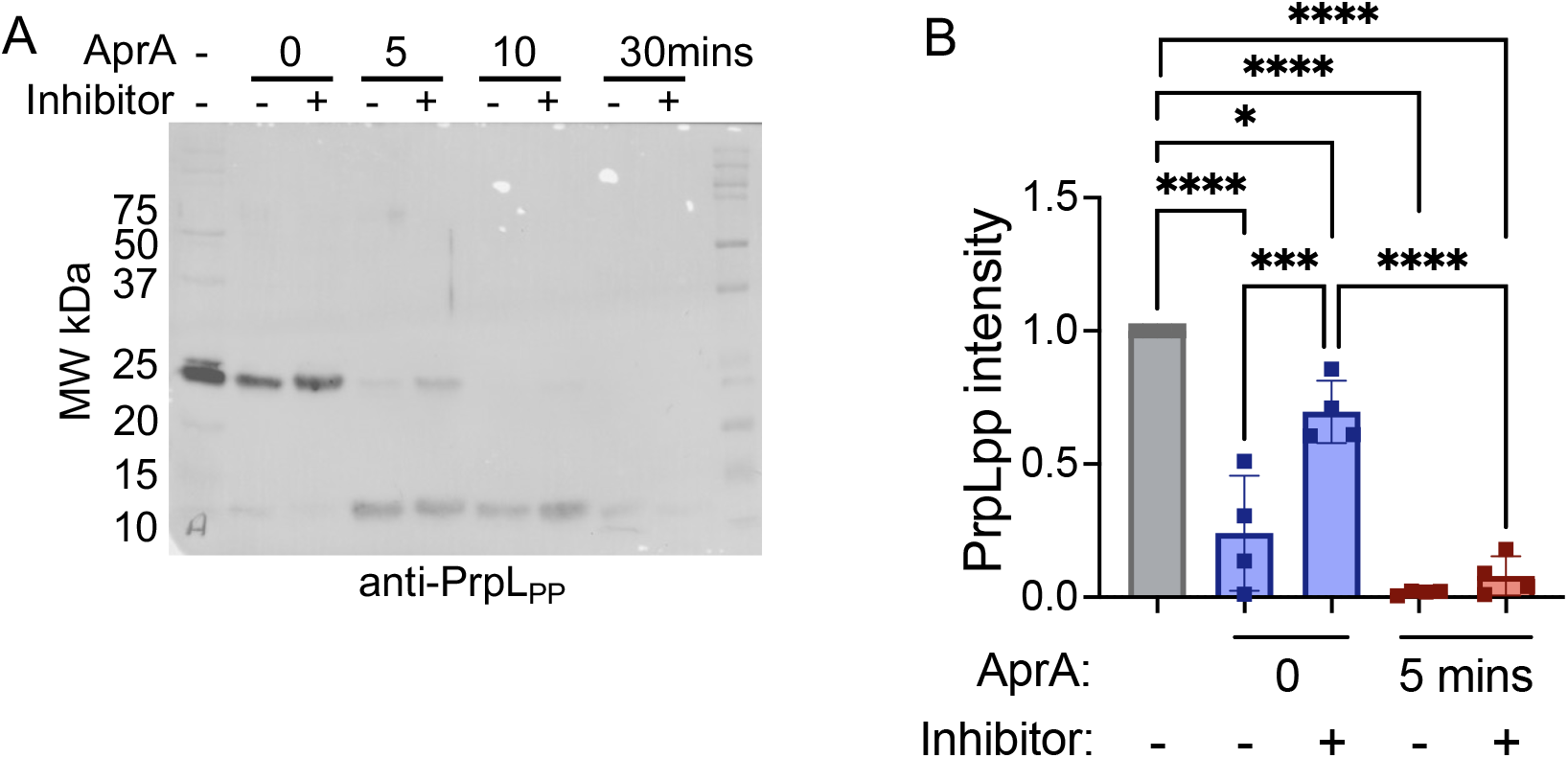
AprA degrades the PrpL propeptide (PrpL_PP_). A) Purified AprA (0.05 µg), pretreated or not with the metalloprotease inhibitor phosphoramidon for 30 min, was incubated with purified PrpL_PP_ (5 µg) at 37 °C for 0, 5, 10, or 30 min. As a control (farthest lane on left), PrpL_PP_ was incubated alone for 30 min under the same conditions. Samples were analyzed by western blotting with anti-PrpL_PP_ antibody. The smaller bands around 10kDa represent intermediate degradation product. B) Quantification of band intensity for the blots (0 and 5 min) were normalized to the no enzyme control (farthest lane on left). Data are represented as mean ± standard deviation and were compared by one-way ANOVA with correction for multiple comparisons. Non-significant = p >0.05, * = p <0.05, *** = p <0.001, and **** = p <0.0001. Comparisons not otherwise indicated are non-significant.

Finally, we investigated whether AprA activates pro-PrpL under native conditions by analyzing the secretome of a *P. aeruginosa ΔaprA-ΔlasB* strain. Cell-free culture samples were incubated with AprA at different concentrations. Immunoblotting with an anti-PrpL antibody showed that AprA was highly effective in processing pro-PrpL to generate mature PrpL, even at very low concentrations (1:400) (Fig. 9A), whereas PrpL_PP_ decreased drastically (Fig. 9A & B). Under the same incubation conditions, LasB treatment of the Δ*aprA*-Δ*lasB* supernatant did not process pro-PrpL to produce significant amounts of mature PrpL until the concentration reached a 1:1 ratio (Fig. 9A & B). Moreover, the PrpL-specific activity in the Δ*aprA*-Δ*lasB* supernatant activated by AprA was much higher than that activated by LasB at lower concentrations (Fig. 9C). It is worth noting that we confirmed that our purified AprA is free of LasB and *vice versa* via western blotting against anti-AprA and anti-LasB antibodies (Supplementary Fig. 1), as well as confirming lack of cross-reactivity using specific enzymatic activity assays. These data suggest that AprA is a much more potent activator of PrpL during its natural maturation process than LasB.

**Figure 9.**
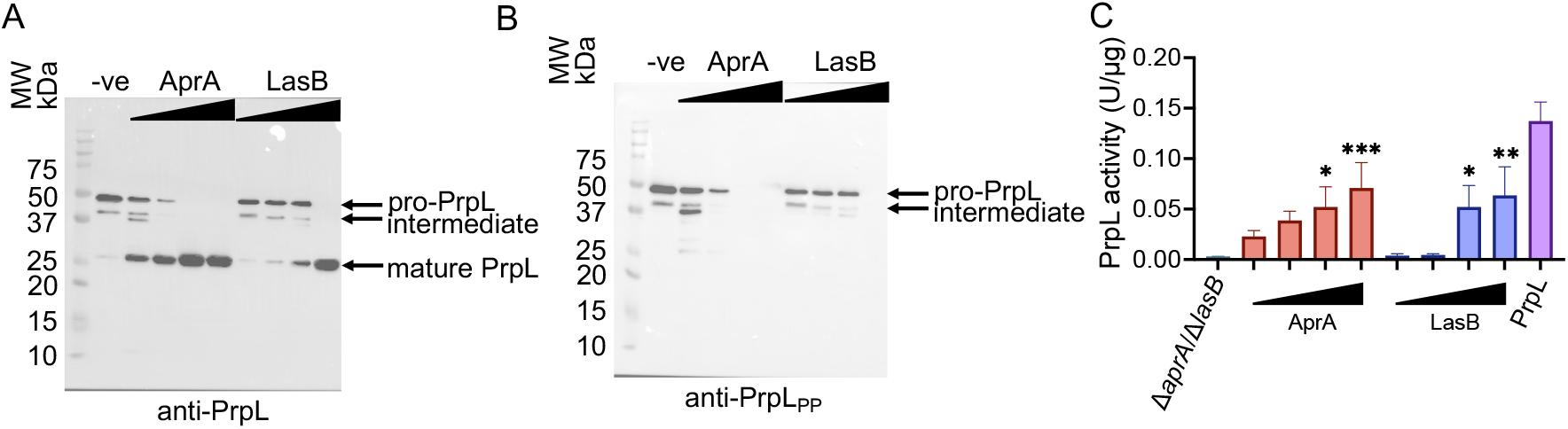
AprA processes pro-PrpL to mature PrpL more efficiently than LasB in the Δ*aprA*Δ*lasB* culture supernatant. Complementation assays were performed by incubating 1 µg of Δ*aprA*Δ*lasB* culture supernatant with increasing concentrations (0.0025, 0.01, 0.1, 1 µg) of purified AprA or LasB or PBS (shown as -ve), followed by western blotting with anti-PrpL (A) and anti-PrpL_PP_ (B) antibodies, and PrpL activity measurement (C). AprA efficiently processed pro-PrpL to the mature form even at the lowest concentration tested (0.0025 µg), whereas LasB required 1 µg for comparable processing. Consistently, PrpL activity was detectable at all AprA concentrations but only at the two highest LasB concentrations. Data are represented as mean ± standard deviation and were compared by one-way ANOVA with correction for multiple comparisons. Non-significant = p >0.05, * = p <0.05, ** = p <0.01, and *** = p <0.001. Comparisons not otherwise indicated are non-significant.

## Discussion

The ability of *P. aeruginosa* to thrive in diverse environments is reflected in its capacity to infect a wide range of hosts (1,2,32–35), highlighting its interactions with varied organisms across ecosystems. This adaptability, driven by shared virulence factors and conserved host immune responses (34), positions *P. aeruginosa* as an ideal model for uncovering evolutionarily conserved mechanisms of innate immune signaling. One example is the type 2-secreted protease PrpL, which has been shown to modulate the immune response to *P. aeruginosa* in several different ways across multiple model systems. In the mealworm *Tenebrio molitor*, PrpL was found to interfere with the production of antimicrobial peptides through disruption of a Toll signaling pathway (36). In *Arabidopsis thaliana*, PrpL was found to activate MPK3 and MPK6 through an immune pathway involving G-protein signaling and the scaffold protein receptor for activated C kinase 1 (RACK1) (20). Moreover, PrpL has been reported to have a higher prevalence among strains isolated from the lungs of CF patients than environmental isolates (37), implicating an important role for PrpL in the pathogenesis of *P. aeruginosa* in the CF lung. PrpL has been identified as a key virulence factor in *P. aeruginosa* eye, skin and lung infections (17,38). However, the detailed impacts of PrpL on inflammatory responses in the mammalian system and the signaling pathways involved are not known.

While an effective immune response to pulmonary *P. aeruginosa* relies on proinflammatory signals to recruit defensive cells, an overly robust or sustained cytokine release can itself become pathogenic, causing systemic inflammation, tissue damage, and mortality (1,2). A well-regulated inflammatory response is therefore crucial, as it balances effective antibacterial defense with the preservation of tissue function (39,40). The transcriptional regulatory networks that achieve this critical balance during *P. aeruginosa* infection, however, remain incompletely elucidated. Our finding that in the mouse lungs purified PrpL robustly activates the transcription factor AP-1 complex and triggers strong proinflammatory cytokine production (Figs. 1 and 6), further confirms its role as a relevant virulence factor in *P. aeruginosa* lung infection and reveals certain transcriptional regulatory pathways that contribute to PrpL-elicited inflammatory responses. Previous studies have reported roles for AP-1 in *P. aeruginosa* lung infections. In particular, *P. aeruginosa* type III-secreted effector ExoU was shown to modulate gene expression in lung epithelial cells partly mediated by activation of the AP-1 transcription factor c-Fos (41). One study demonstrated that both AP-1 and NF-κB control the induction of human β-defensin-2 in keratinocytes during exposure to *P. aeruginosa* and IL-1β (42). Our study provides strong evidence that AP-1 transcriptional control contributes to PrpL-induced inflammation.

In support of this, both the catalytically inactive PrpL variant (PrpL^S198A^) and the use of a dipeptide inhibitor of PrpL activity (LasBi) led to a loss of AP-1 activation (Figs. 1 & 6). Consequently, downstream inflammatory responses, including cytokine production and clinical symptoms, were also lacking in mouse lungs when PrpL activity was absent (Figs. 1 & 6). It is worth noting that phospho-c-Fos, but not phospho-c-Jun, levels were significantly different between groups, although there was a trend toward lower phospho-c-Jun levels in the LasBi-treated group compared to untreated PrpL and in the PrpL^S198A^ mutant compared with WT PrpL (Figs. 1 and 6). AP-1 is a heterogeneous transcription factor complex that can be assembled from multiple Jun and Fos family members (43). Therefore, it is possible that phospho-c-Fos forms homodimers or partners with an alternative AP-1 component that was not examined in the present study. Taken together, the robust differences observed in phospho-c-Fos strongly support AP-1 activation by PrpL activity, even in the absence of significant changes in phospho-c-Jun. These findings suggest that c-Fos may play a more prominent role than c-Jun in mediating the AP-1 response induced by PrpL in our experimental system.

The observation that PrpL activity is important for activation of mammalian inflammatory responses is also consistent with our previous work that identified a unique requirement of PrpL enzymatic activity for activating innate immunity in plants, unlike the canonical MAMP-triggered innate immune responses that only require a conserved epitope (20). Critically, our findings that PrpL’s enzymatic activity is required to elicit proinflammatory responses in mouse lungs provides a rationale for targeting it therapeutically to alleviate the excessive inflammation and lung tissue damage caused by *P. aeruginosa* lung infections.

Protease inhibitors hold potential as alternative antimicrobial therapies, aiming to attenuate virulence without promoting antibiotic resistance (44). Inhibition of PrpL, such as by its own propeptide PrpL_PP_, has been shown to attenuate infections in invertebrates and mice, including skin and lung infections (38). However, the 19-kDa full-length propeptide is not ideal for drug development because of its potential high immunogenicity. In order to understand how PrpL_PP_ inhibits PrpL and aid our search for a potent inhibitory peptide with minimum length, we used X-ray crystallographic analysis to solve the co-structure of PrpL bound to PrpL_PP_ (Fig. 5). Surprisingly, PrpL caused the disappearance of the PrpL_PP_ when incubated long-term for crystallization, and only the addition of the protease inhibitor TLCK prevented the degradation of the propeptide in the PrpL^S198A^-PrpL_PP_ complex. Although the co-structure revealed an 8-amino acid peptide fragment from PrpL_PP_ that blocks the catalytic site of PrpL, providing the potential inhibitory mechanism, the synthetic octamer did not show any inhibition of PrpL activity. This is similar to a previous study of the endogenous AprA inhibitor AprI in *P. aeruginosa*, which showed that truncated AprI lost its inhibitory effect (45). PrpL_PP_ binds to PrpL through extensive interactions and the synthetic shortened peptides might not bind properly or only weakly to their target proteases.

To circumvent the lack of activity of the 8-amino acid peptide, structure-guided selection of a previously studied dipeptide inhibitor, LasBi, demonstrated dose-dependent inhibition of PrpL activity *in vitro* and effectively abrogated PrpL-induced inflammatory responses *in vivo*, including decreased transcription factor AP-1 activation, drastically reduced proinflammatory cytokine production, and alleviated clinical symptoms (Fig. 6). LasBi is a relatively weak inhibitor of PrpL, which is not entirely unexpected given that its sequence is only modestly similar to that of the natural propeptide near the active site. Nevertheless, this work establishes proof-of-principle that pharmacological inhibition of PrpL activity and virulence using synthetic dipeptides is feasible. With further optimization to enhance potency, such inhibitors hold potential as therapeutic candidates for mitigating *P. aeruginosa*-induced lung inflammation and tissue damage. Targeting the virulence factor PrpL avoids exerting direct selection pressure on bacteria, thereby potentially reducing the likelihood of developing drug resistance, a significant issue with treatments for drug resistant *P. aeruginosa* infections. In addition, targeting secreted PrpL will bypass the membrane permeability and efflux pump concerns related to the development of anti-bacterial compounds targeting Gram-negative bacteria.

The identification of PrpL as a potent inducer of host inflammatory signaling represents an important advance in our understanding of host-pathogen interactions. Our findings also highlight the complexity of inflammation during *P. aeruginosa* infection. Although PrpL exhibited a robust and biologically significant pro-inflammatory role when instilled directly in the experimental systems examined, mice infected with the Δ*prpL* mutant did not show significantly reduced inflammatory responses compared with those infected with WT bacteria. This is likely due to the substantial functional redundancy among the numerous inflammatory effectors produced by *P. aeruginosa*, which may compensate for the loss of a single factor *in vivo*. Given this redundancy in the infection model used in the current study, targeting PrpL as a therapeutic strategy to dampen inflammation during *P. aeruginosa* lung infection would likely need to be combined with approaches that simultaneously target other pro-inflammatory bacterial factors. However, the Δ*prpL* mutant has demonstrated attenuated virulence in other infection models, particularly in corneal infection models. PrpL may represent a promising anti-virulence therapeutic target in immune-privileged sites such as the cornea, although its inhibition alone may not be sufficient to specifically or completely suppress host inflammatory responses at sites such as the lung. Future work can use structural information in combination with medicinal chemistry to further develop other short peptides or small molecule inhibitors that have greater potency and attractive pharmaceutical properties.

Extracellular proteases are activated through a hierarchical cascade. Specifically, LasB degrades PrpL_PP_, releasing its active form and then both LasB and PrpL subsequently activate LasA by degrading LasA propeptide (22,46). Although this model has greatly expanded our understanding of sequential protease maturation post-secretion, our findings in this study show that LasB and another secreted protease AprA can individually activate PrpL by degrading PrpL_PP_ (Figs. 7-9). It is worth noting that LasB and AprA exhibit cooperative roles in degrading other substrates, including bacterial flagellin (29,30) and host glycoprotein mucin enriched in CF lungs (47), highlighting their functional redundancy. This cooperation underscores the importance of proteolytic activity in *P. aeruginosa* virulence and its ability to modulate immune signaling as well as mucin-based nutrient acquisition and host environment adaptation. Notably, both AprA and LasB are zinc metalloproteases and metalloprotease inhibitors have been extensively investigated in clinical trials in various cancers and inflammatory diseases (48). The extensive preexisting clinical data on these inhibitors can facilitate their potential repurposing as a new antibacterial therapy for inhibiting lung damage caused by bacterial proteases during infections.

## Materials and Methods

### Bacterial strains, plasmids, and culture conditions

All the bacterial strains, plasmids, and cloning primers used in this study are listed in Table 2. *P. aeruginosa* and *E. coli* strains were grown overnight (18h) at 37 °C in Luria-Bertani or M9 medium with shaking at 200 rpm. Agar was added to the media at 1.5% (w/v) to prepare solid media. For the cultivation of plasmid-carrying *E. coli*, the antibiotics ampicillin 100 μg/ml and kanamycin 50 μg/ml were used, and 0.4 mM IPTG (isopropyl-1-thio-β-D-galactopyranoside) was added to the culture media to induce protein expression. For *P. aeruginosa,* carbenicillin 300 μg/ml or kanamycin 300 μg/ml was used for plasmid selection. PAO1/*ΔprpL* mutant was created and confirmed according to the published protocol (20).

**Table 2.**
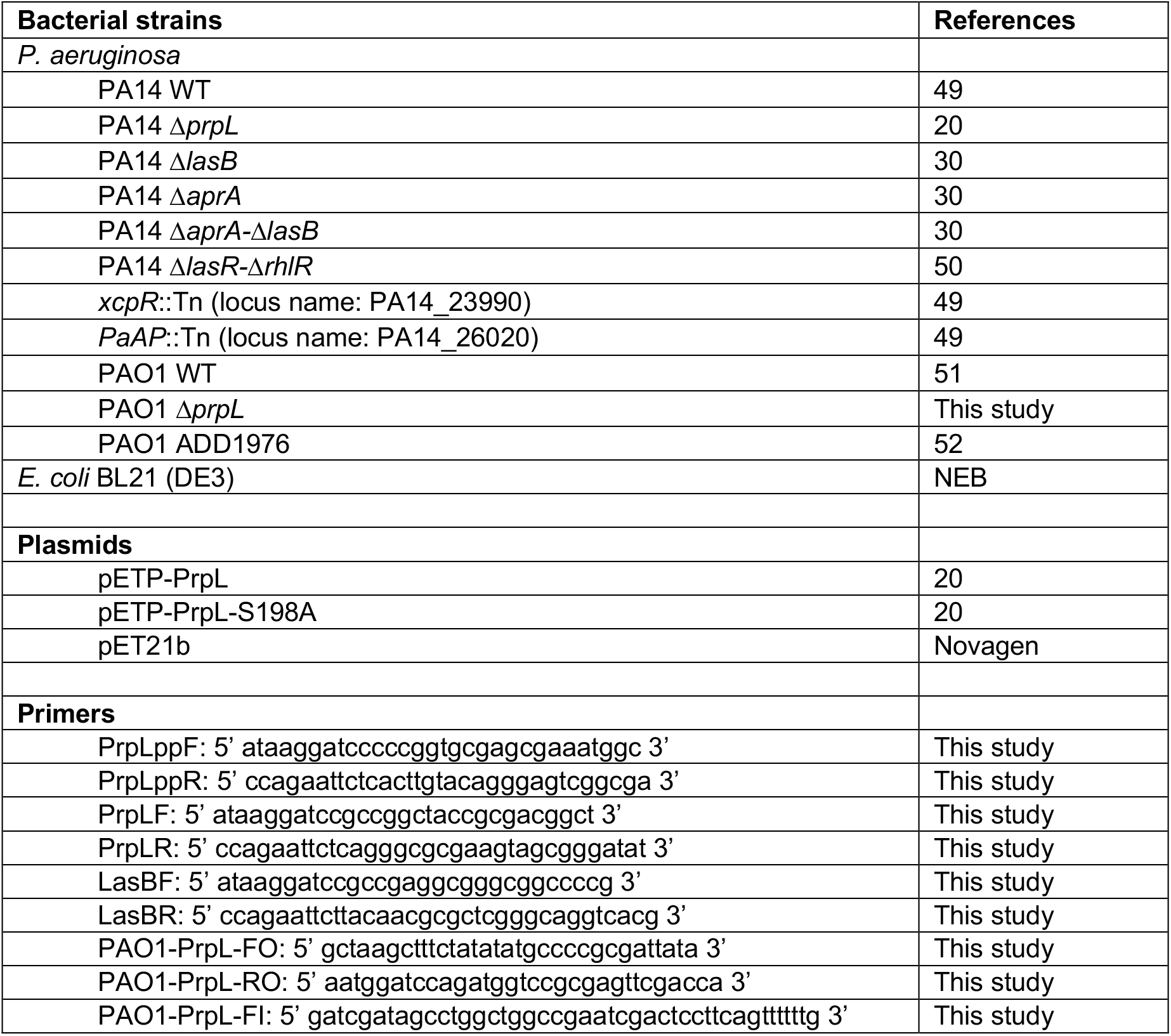
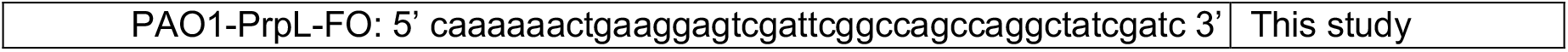
Bacterial strains, plasmids and cloning primers used in this study.

### Culture supernatant preparation from *P. aeruginosa* strains

For the preparation of culture supernatants, WT and mutant *P. aeruginosa* cells were grown in 5 ml of LB with 200 rpm shaking at 37 °C overnight. The next morning, optical density at 600 nm was adjusted and cells were diluted 1:100 into fresh 10 ml of M9 broth. After further cultivation with 200 rpm shaking at 37 °C for 24 hours, cells were removed by centrifugation at 6,000 × *g* for 10 min at 4 °C followed by filtration through 0.22 μm syringe filter (Satorious). The collected supernatants were concentrated 20 times through 10 kDa cut-off centricon (Amicon® Ultra Centrifugal Filters), and the total protein in each sample was quantified using a BCA protein kit (Pierce™ BCA Protein Assay Kit).

### Protein purification, antibody production, and peptide synthesis

*PrpL_PP_* sequence was amplified from *P. aeruginosa* genomic DNA using primers and cloned into a pET21b expression plasmid modified to include an N-terminal 6xhistidine tag and tobacco etch virus protease recognition sequence (pET21b-H-TEV-PrpL_PP_). The H-TEV-PrpL_PP_ was purified in lysis buffer (20 mM Tris-HCl pH 8, 100 mM NaCl, 5% glycerol) by applying cell lysate to Ni-NTA resin (IMAC Sepharose 6 Fast Flow; Cytiva). TEV-protease (0.2 mg per liter of cell culture) was added to the elution and the mixture was dialyzed overnight against lysis buffer. The sample was then passed through Ni-NTA resin and PrpL_PP_ was isolated by collecting the flowthrough. The recombinant PrpL_PP_ protein was used for the degradation assay, crystallographic analysis, and polyclonal antibody production. Recombinant PrpL, PrpL_PP_, and LasB were used for polyclonal antibody production only, whereas endogenous PrpL, LasB, and AprA were purified from *P. aeruginosa* supernatants for the degradation assay and crystallographic analysis.

PrpL and the catalytic mutant PrpL^S198A^ were purified from *P. aeruginosa* supernatant as described previously (20). AprA and LasB were also purified from the *P. aeruginosa* PA14 supernatant as described previously, with the exception that the supernatant of the transposon mutant of the aminopeptidase (locus name: PA14_26020) was used for the AprA purification to separate the aminopeptidase and AprA that have similar molecular weights (53). All proteins were further purified by size exclusion chromatography with HiLoad 16/60 Superdex 200 gel filtration column (Cytiva). Endotoxin was removed from purified PrpL and PrpL^S198A^ using Pierce™ High Capacity Endotoxin Removal Spin Columns (Thermo Scientific) following the manufacturer’s protocol, and endotoxin removal was verified using a ToxinSensor™ Chromogenic LAL Endotoxin Assay (GenScript). The purified proteins were filter-sterilized using the Ultrafree-CL Centrifugal Filter Devices (Millpore), aliquoted, and stored at −80 °C until further use. Purity of the proteases was assessed by electrophoresis on 12.0% SDS-PAGE gels followed by Coomassie blue staining and western blot using anti-PrpL, anti-AprA, and anti-LasB antibodies. The polyclonal anti-PrpLpp, anti-PrpL, and anti-LasB antibodies were raised in New Zealand White rabbits and purified by protein A from antiserum (Vivitide). The anti-AprA antibody was produced as described previously (54). The small peptides, including LasBi (mercaptoacetyl-Phe-Tyr-NH_2_) (28,55), were synthesized on a 0.5 mmol scale (56) and the octamer derived from the PrpL_PP_ (Ac-Lys-Pro-Gly-Thr-Pro-Leu-Gln-Val-NH_2_) was synthesized on a 0.1 mmol scale using manual solid phase peptide synthesis. Peptides were synthesized on Rink Amide AM resin using commercially available Fmoc-protected L-amino acids. Standard peptide synthesis procedures were employed using O-(1H-6-Chlorobenzotriazole-1-yl)-1,1,3,3-tetramethyluronium hexafluorophosphate (HCTU) as the coupling reagent. Crude peptides were purified using preparatory high performance liquid chromatography (HPLC) resulting in >95% purity. Isolated peptides were characterized using analytical reverse-phase HPLC linked to electrospray mass spectrometry (MS).

### Protease activity assays, inhibition, and PrpL_PP_ degradation

The chromogenic substrate chromozym PL (N-(p-tosyl)-Gly-Pro-Lys-4-nitroanilide acetate salt, Sigma) was used as a substrate for PrpL protease as described previously (8). After dissolving substrate by 2 mg/ml in glycine-tween 20 buffer (100 mM glycine, 0.2%Tween 20, pH 8.0), 20 µL was mixed with 1 µg purified PrpL or 5 µg concentrated total proteins from *P. aeruginosa* culture supernatant in 150 mM NaCl in 50 mM Tris (pH 8.0) with a total reaction volume of 100 µL. The reaction mixture was immediately placed into a microtiter plate reader and the optical density at 410 nm was read at 1-min intervals for 20 min. The units of enzyme were calculated from the change in optical density (ΔA) per min. One unit of PrpL activity was defined as ΔA/min × total assay volume/sample volume × E@410 nm × light path (cm), where the extinction coefficient (E) of the product (Paranitroanilide) at 410 nm was 9.75 and the light path was 0.53 cm under the assay conditions (8,57). AprA activity was quantified using the substrate Hide-Remazol Brilliant Blue R (Sigma) as described previously (58). LasB activity was determined using an elastin-Congo red method, as previously described (46). These specific enzymatic assays were also used to examine the purity of all purified proteases and no cross-reactivity was observed in purified LasB, AprA, or PrpL.

The serine protease inhibitor TLCK was used at 2 mM in the crystallization of the PrpL^S198A^-PrpL_PP_ complex. The synthetic octamer was used at final concentrations of 1 µM, 10 µM, and 100 µM in the PrpL activity assay, and LasBi was used at 1 µM, 10 µM, 100 µM, 200 µM, 250 µM, 500 µM and 1 mM. Both peptide inhibitors were incubated with PrpL at 37 °C for 15 minutes prior to addition of the substrate. The metalloprotease inhibitor was used at 40 μM to be incubated with AprA for 30 minutes at 37 °C in the PrpL_PP_ degradation assay.

Purified proteases were mixed with 5 μg purified PrpL_PP_ or 1 μg of supernatant proteins from the PA14 *ΔaprA-ΔlasB* double mutant at the w/w ratio of 1:1, 1:10. 1:100, and 1:400 for different time points from 0 to 30 minutes at 37 °C with or without specific inhibitors. The reaction products were separated on 12.0% SDS-PAGE gels followed by western blot using anti-PrpL and anti-PrpL_PP_ antibodies to examine the PrpL_PP_ degradation dynamics.

### X-ray crystallographic analysis of the PrpL^S198A^-PrpL_PP_ complex

The PrpL_PP_-PrpL^S198A^ complex was isolated by gel filtration in 20 mM Tris pH 8.0, 100 mM NaCl, 5% glycerol, and 50 μM TLCK. The complex was concentrated to 2 mg/mL then screened via microbatch under paraffin oil using the JBScreen Classic HTS I/II kit (Jena Bioscience). Crystal hits were optimized, and final crystallization conditions were obtained by mixing equal volumes of 4.2 mg/mL protein with 100 mM MES pH 6.5, 2.8 M ammonium sulfate and 5% (w/v) MPD. Diffraction data were collected at the Canada Light Source (CMCF-ID beamline) and processed using the associated MXProc software. The structure was solved using Phenix (59) through molecular replacement using the AlphaFold2-predicted structures (60) of PrpL and PrpL_PP_ as search models. A final model was refined to 3.3 Å using Phenix and manual building with COOT (61). Final coordinates were deposited to the protein data bank (62) (accession number 9OMD).

### Intratracheal *P. aeruginosa* infection and PrpL instillation

C57BL/6 mice were originally sourced from the Jackson Laboratory (Bar Harbor, ME) and bred at the Carleton Animal Care Facility at Dalhousie University. Mice used in experiments were 8-12 weeks of age. Approximately equal numbers of male and female mice were used for each experiment. The experimental protocols (23-073 and 25-084) were approved by the University Committee on Laboratory Animals based on adherence to Canadian Council on Animal Care guidelines. For *P. aeruginosa* infection experiments, an otoscope was used to visualize the tracheal opening, and 40 μL of PA14 or PA14/Δ*prpL* or PAO1 with its corresponding PAO1/Δ*prpL* inoculum at the dose of 5×10^5^ and 1×10^6^ CFU in PBS respectively was deposited directly in front of the vocal folds of the tracheal opening using a micropipette. Control mice were inoculated with an equivalent volume of sterile PBS. For direct instillation experiments, purified PrpL, PrpL^S198A^, LasBi-inhibited PrpL, or *P. aeruginosa* LPS (Sigma-Aldrich) were instilled intratracheally using the method described above. Per gram body weight, 100 or 500 ng of PrpL or PrpL^S198A^ or 5 µg of LPS was used, delivered in PBS in a final volume of 50 µL. For LasBi-inhibited PrpL, LasBi was added to PrpL at a final concentration of 200 µM and incubated at 37 °C for 1 hour prior to instillation. Control mice were inoculated with 50 µL of PBS or 200 µM LasBi in PBS.

For both infection and instillation experiments, mice were observed for signs of morbidity and provided supportive care. Weight loss greater than 15% of initial weight or clinical score ≥ 12 were criteria for immediate euthanasia. No mice met these criteria throughout the study period. Mouse weights and clinical scores (assigned using criteria in Supplementary Table 1) were recorded at 4 and 24-hours post infection or instillation.

At 24 hpi mice were anesthetized with inhaled isoflurane before being sacrificed via open cardiac puncture. From infection experiments, BALF was collected for ELISA and lungs were collected for ELISA, western blotting, flow cytometry, and histological assessment of tissue damage using H&E staining. From instillation experiments, BALF was collected for ELISA and lungs were collected for ELISA and western blotting. ELISAs were performed using DuoSet ELISA kits for mouse IL-1β, IL-6, and TNF (R&D Systems) according to the manufacturer’s instructions. For western blotting, antibodies of rabbit origin against mouse IκB, c-Fos, and c-Jun targets were purchased from Cell Signaling Technologies (Supplementary Table 2). Western blot quantification was performed using Image Lab version 6.1 (Bio-Rad). For all targets, each lane’s adjusted volume (intensity) was normalized to GAPDH intensity for the same lane. For flow cytometry analysis, whole lungs were mechanically separated and digested in RPMI-1640 with collagenase D and DNase I for 1 h at 37 °C, filtered through 70 μm strainers, and red blood cells were removed using ammonium-chloride-potassium (ACK) lysis buffer. Cells were washed, resuspended in 2% FBS/PBS, and counted. BALF cells were washed and resuspended in 2% FBS/PBS. Lung and BALF cells were stained with Fixable Viability Dye (eFluor780, eBioscience), and a cell surface receptor antibody panel (Supplementary Table 3) in the presence of TruStain FcX™ PLUS Fc block (Biolegend), to identify macrophage, neutrophil, and dendritic cell subsets. Stained cells were fixed in 2% paraformaldehyde, washed, and stored at 4 °C. Data were acquired on a BD LSR Fortessa with FACSDiva software and analyzed using FlowJo v10.6. For histology, lungs were collected and fixed in 10% neutral-buffered formalin (NBF) for 24-hours, and then washed and stored in 70% ethanol until processing. Lungs were embedded in paraffin, sectioned, and stained with hematoxylin and eosin (H&E). An Axio Observer inverted light microscope was used to take representative images of histology slides. Histology slides were analyzed blindly by a pathologist.

## Statistical analysis

Data analysis and statistical tests were performed using GraphPad Prism (version 10.5.0). Data are expressed as mean ± standard deviation (SD). Statistical significance was set at p<0.05. Outliers were identified using GraphPad’s ROUT method and excluded. Data was tested for normality using the Shapiro-Wilk test (data with p>0.05 was considered normal). Data that met the assumption of normality was analyzed using a one-way or two-way analysis of variance (ANOVA) with a post-hoc Tukey’s multiple comparisons test to compare groups, with alpha error rate at 0.05. Data that did not meet the assumption of normality was analyzed using a non-parametric Kruskal-Wallis test with Dunn’s post-hoc test to compare groups. Weight loss and clinical score data were compared using Mann-Whitney tests due to uneven sample sizes between groups.

## Acknowledgments

We would like to thank Dr. Susanne Häußler at the Helmholtz Centre for Infection Research, Germany for providing the *P. aeruginosa* mutants, Dr. Bart Bardoel at the University Medical Center Utrecht, Netherlands for generously sharing anti-AprA antiserum, and Dr. Frederick M. Ausubel at Massachusetts General Hospital, Boston Massachusetts, for helpful discussions and for reviewing the manuscript. We appreciate the support from Dalhousie University’s Flow Cytometry, Histology and Animal Care facilities. This research was supported by the Canadian Institute of Health Research (CIHR) Project Grant (PJT165970) and the Natural Sciences and Engineering Research Council of Canada (NSERC) Discovery Grant (RGPIN-2023-05928) to Z.C.; the NSERC Discovery Grant (RGPIN-2024-06440) to D.N.L.; the NSERC Discovery Grant (RGPIN-2023-05922) and Canada Foundation for Innovation (CFI) John R. Evans Leaders Fund (JELF44543) to C.L.C.; and the National Natural Science Foundation of China grant (81801962) and the President Foundation of Nan Fang Hospital, Southern Medical University grant (2021B019) to X.Z. X-ray crystallography studies were performed using beamline CMCF-ID at the Canadian Light Source, a national research facility of the University of Saskatchewan, which is supported by the CFI, NSERC, National Research Council (NRC), CIHR, the Government of Saskatchewan, and the University of Saskatchewan. We also thank Dr. Michel Fodje for support in diffraction data processing. S.M.D. received the Dalhousie Faculty of Medicine I3V Research Associate Award; L.D.B. is a recipient of the Canada Graduate Scholarship (CGS) Masters Award, Nova Scotia Graduate Scholarship (NSGS), and a CIHR CAN-AMR-Net Health Research Training Program scholarship; T.N. and R.N. were partially supported by Dalhousie Faculty of Medicine Graduate scholarships; R.N. also received an NSGS; S.L.G. received CGS Doctoral Award and NSGS.

