## Supplementary figures and tables for "Inhibition of *Pseudomonas aeruginosa-*secreted protease IV reduces lung inflammation"

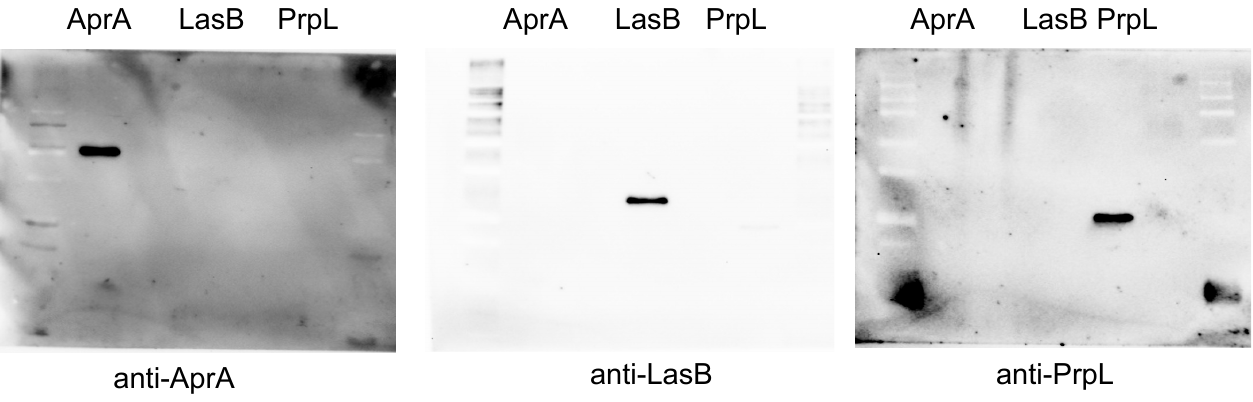

**Supplementary Figure 1. The proteases purified from the native *P. aeruginosa* supernatants were free of other proteases.** The purity of AprA, LasB and PrpL used in this study were tested by western blot using anti-PrpL, anti-AprA, and anti-LasB antibodies. The target protease only reacts with their corresponding antibodies, but not the other two.

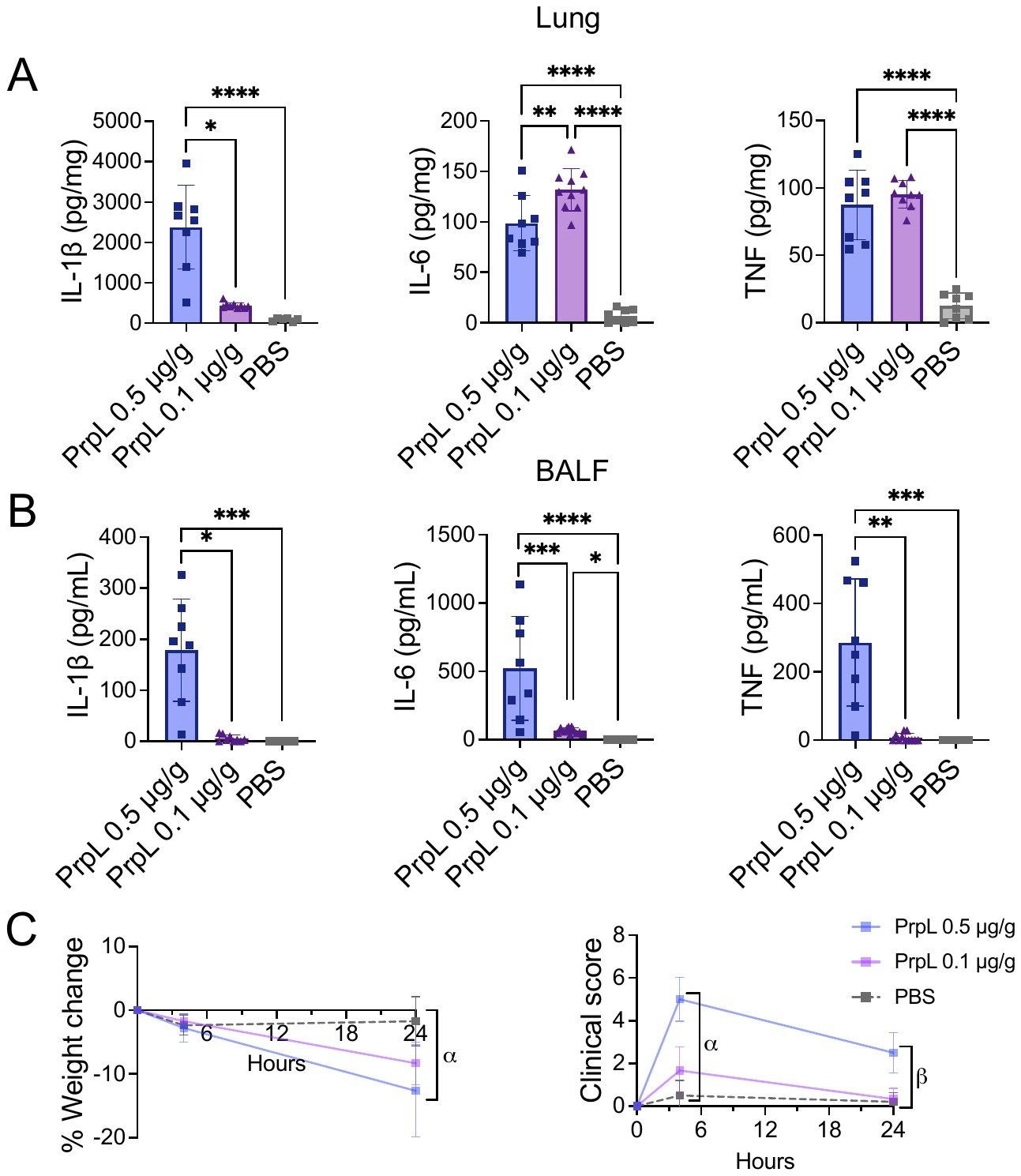

**Supplementary Figure 2. Instillation of low dose (0.1 µg/g) PrpL induces moderate clinical symptoms and inflammatory cytokine levels.** Purified PrpL was instilled into the lungs of mice at a dose of 0.1 µg/g body weight. Lungs and bronchoalveolar lavage fluid (BALF) were collected 24-hours post-instillation. Data is plotted alongside data from 0.5 µg/g PrpL instillation data and PBS control data from Figure 1. A) Concentrations of inflammatory cytokines IL-1β, IL-6, and TNF in lung homogenate, expressed per mg total lung protein. C) Concentrations of IL-1β, IL-6, and TNF in BALF, expressed per mL BALF. D) Percentage change in mouse body weight measured at 4- and 24-hours post-instillation of PrpL, relative to body weight pre-instillation. α represents p <0.0001 (****) for all comparisons. Clinical scores assigned at 4- and 24-hours post-instillation. α represents the following comparisons: PrpL 0.5 µg/g vs PrpL 0.1 µg/g = ****; PrpL 0.5 µg/g vs PBS = ****; and β represents the following: PrpL 0.5 µg/g vs PrpL 0.1 µg/g = ***; PrpL 0.5 µg/g vs PBS = ****. Data are represented as mean ± standard deviation and were compared by one-way ANOVA with correction for multiple comparisons or Mann-Whitney test where applicable. Non-significant = p >0.05, * = p <0.05, ** = p <0.01, *** = p <0.001, and **** = p <0.0001. Comparisons not otherwise indicated are non-significant. *N* = 3 for 0.1 µg/g PrpL males, & *N* = 6 for 0.1 µg/g PrpL females.

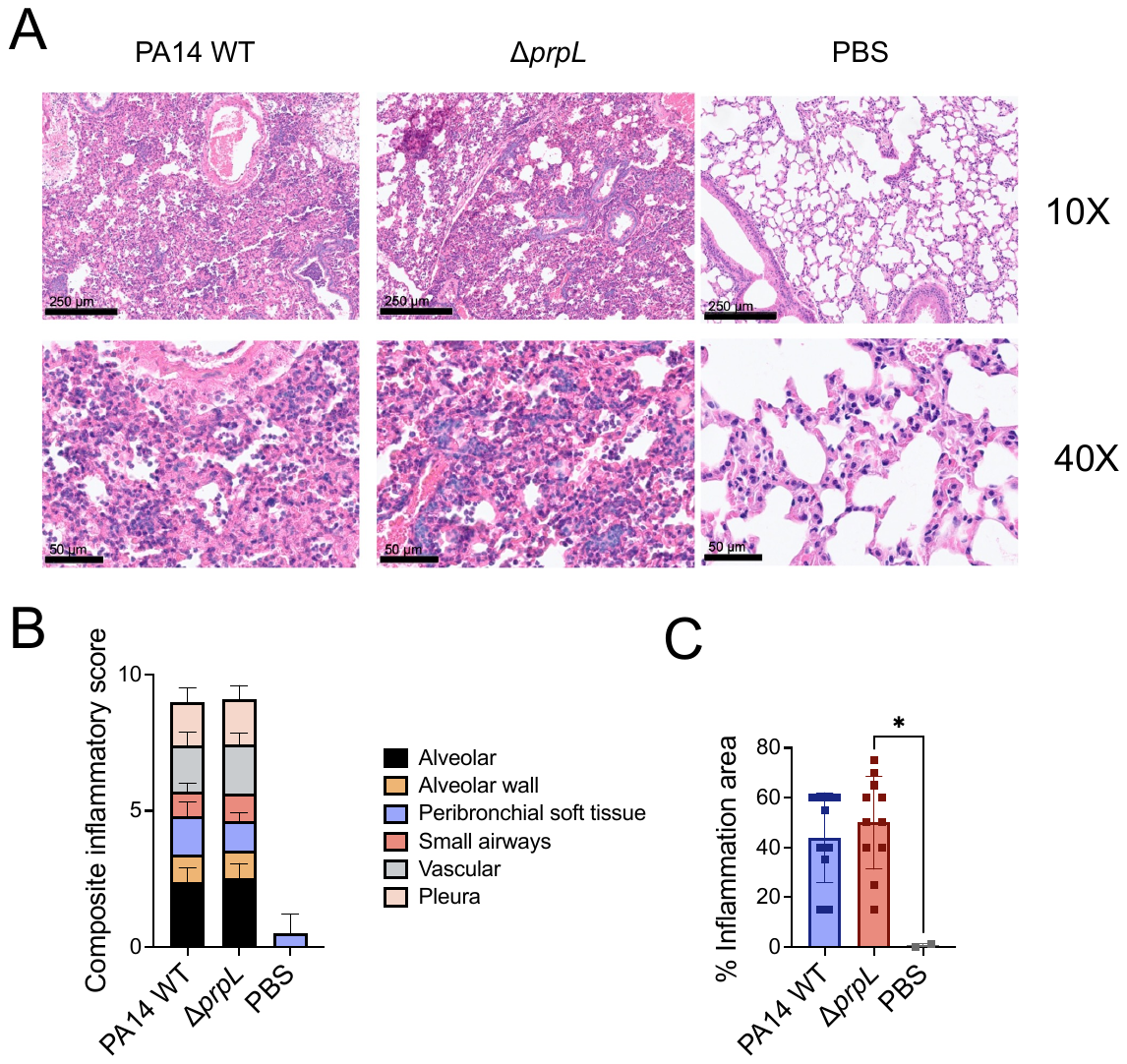

**Supplementary Figure 3. Δ*prpL* and PA14 induce similar levels of histological lung injury *in vivo*.** Mice were intratracheally infected with 5x10^5^ CFU of *P. aeruginosa* PA14 wild-type (WT) or Δ*prpL* mutant. Lungs were collected 24-hours post-infection. A) Representative microscopy images (top: 10X, scale bar 250 μm; bottom: 40X, scale bar 50 μm) of prepared H&E-stained lung sections for control (PBS), PA14 WT-infected, and Δ*prpL*-infected mice. B) Composite inflammatory score, representing the sum of inflammatory scores assigned to individual areas of the lung. Lung areas were scored on a scale of 0 to 3, with 0 indicating a lack of inflammation and 3 representing most severe inflammation. C) Percentage area of inflammation was estimated by examining complete cross-sections of the lungs and scored in 5% increments. Data are represented as mean ± standard deviation and were compared by one-way ANOVA with correction for multiple comparisons. Non-significant, p >0.05, * = p <0.05. *n* ≥10 per group for each sex, except *n* ≥6 for PBS controls.

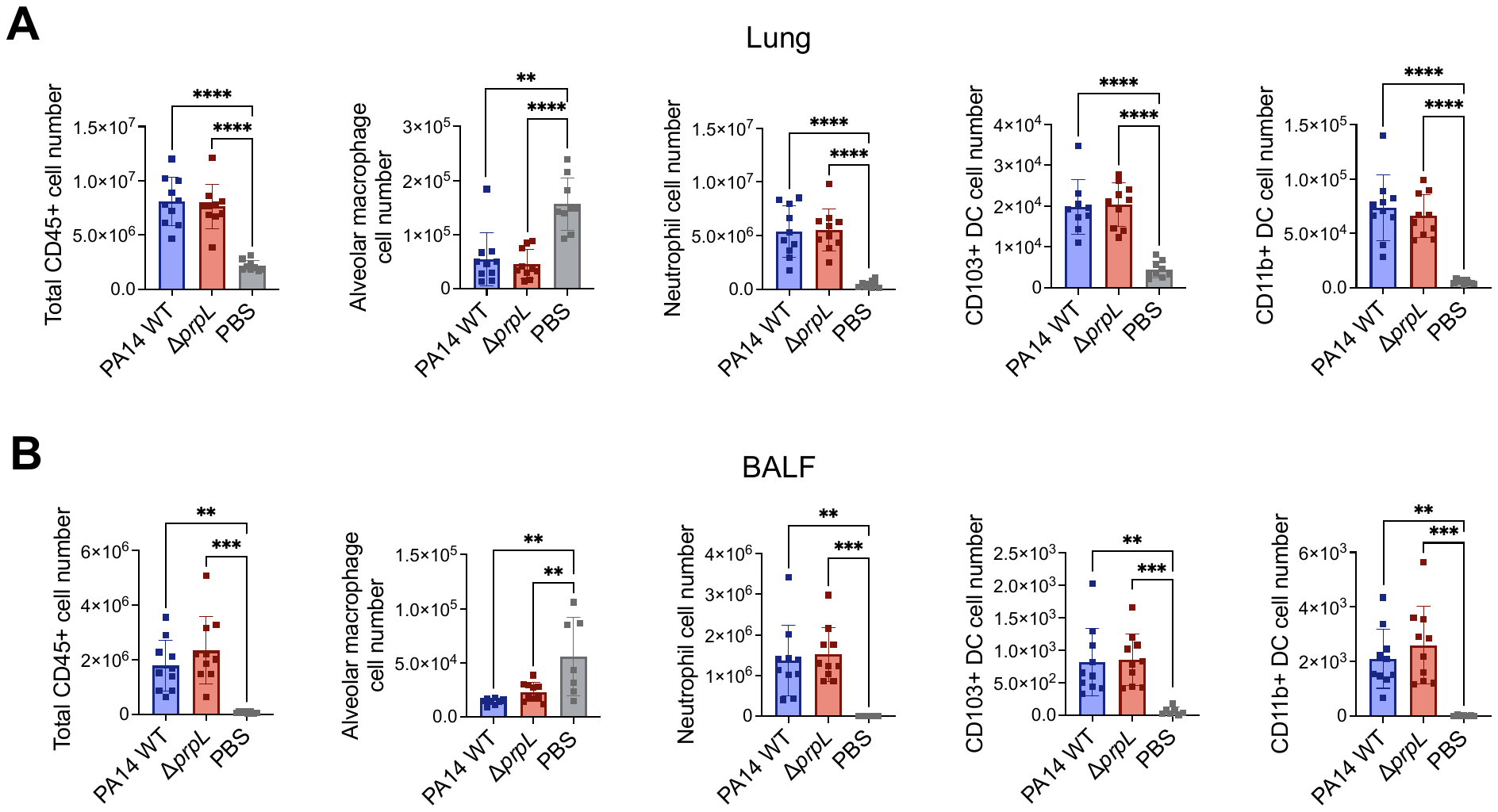

**Supplementary Figure 4. Immune cell recruitment does not differ significantly between Δ*prpL* and PA14 WT *in vivo*.** Mice were intratracheally infected with 5x10^5^ CFU of *P. aeruginosa* PA14 wild-type (WT) or Δ*prpL*. CD45+ cells were isolated from lungs and BALF collected 24-hours post-infection and analyzed using flow cytometry. A) Cell number for lung immune cell populations, calculated by multiplying the percentage of CD45+ cells by the total number of the cells within the samples, which were enumerated using a hemocytometer. B) Cell number for BALF immune cell populations. Data are represented as mean ± standard deviation and were compared by one-way ANOVA with correction for multiple comparisons. Non-significant, p >0.05, ** = p <0.01, *** = p <0.001, and **** = p <0.0001. *n* ≥10 per group, except *n* ≥6 for PBS controls.

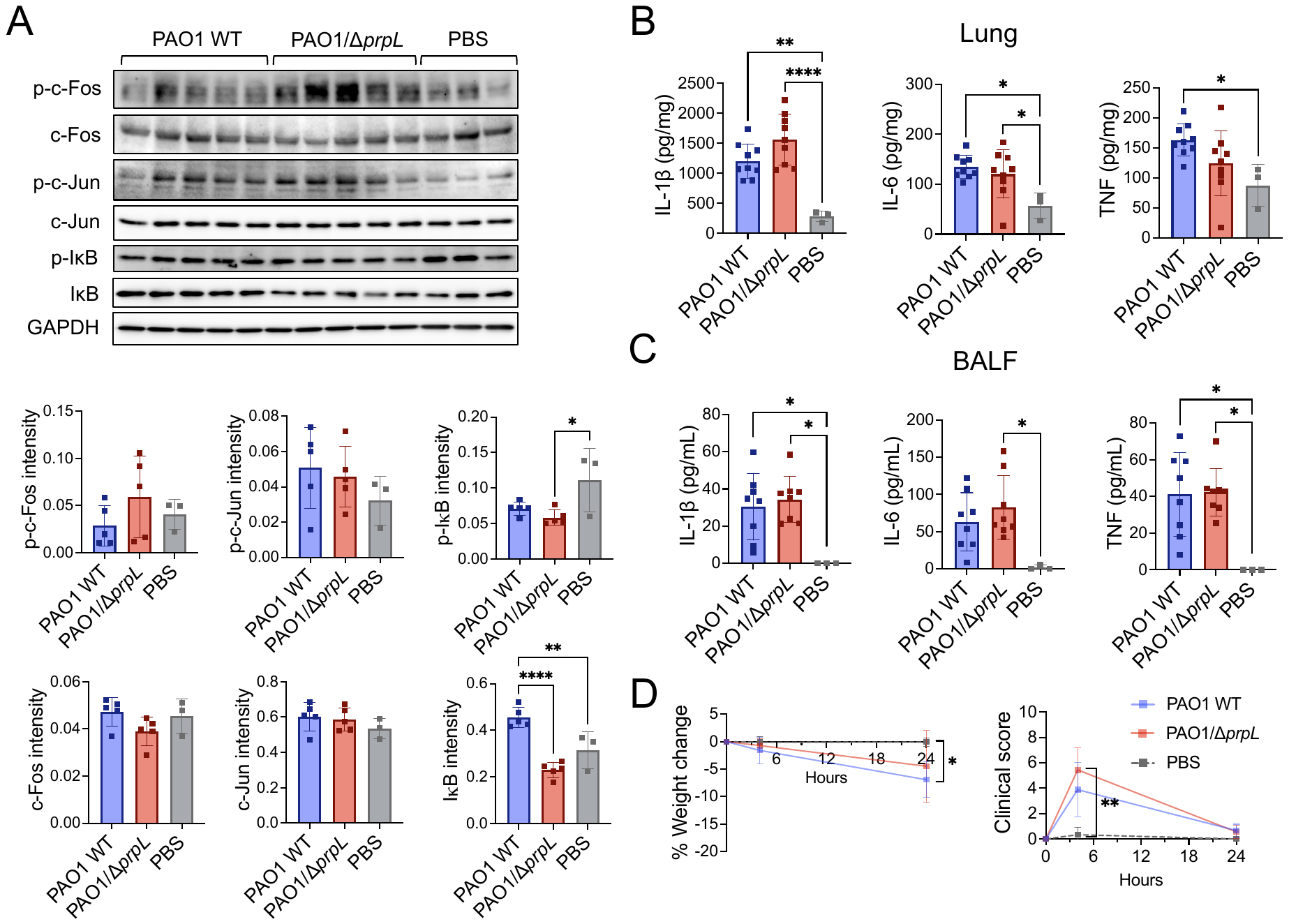

**Supplementary Figure 5. Comparisons of inflammatory responses in mice infected with P. aeruginosa WT PAO1 or PAO1/Δ*prpL* mutant.** Mice were intratracheally infected with 1x10^6^ CFU of *P. aeruginosa* PAO1 wild-type (WT) or PAO1/Δ*prpL*. Lungs and BALF were collected 24-hours post-infection. A) Western blots detecting total and phosphorylated (p-) c-Fos, c-Jun, as well as IκB in lung homogenate. Quantification of band intensity for the blots were normalized to GAPDH loading control. Each lane represents a lung homogenate sample from an individual mouse, with different lanes corresponding to different mice. B) Concentrations of inflammatory cytokines IL-1β, IL-6, and TNF in lung homogenate, expressed per mg total lung protein. C) Concentrations of IL-1β, IL-6, and TNF in BALF, expressed per mL BALF. D) Percentage change in mouse body weight measured at 4- and 24-hours post-infection. Clinical scores assigned at 4- and 24-hours post-infection. Data are represented as mean ± standard deviation and were compared by one-way ANOVA with correction for multiple comparisons or Mann-Whitney test where applicable. Non-significant, p >0.05, ** = p <0.01, *** = p <0.001, and **** = p <0.0001. Comparisons not otherwise indicated are non-significant. *n* ≥9 per group, except *n* ≥6 for PBS controls.

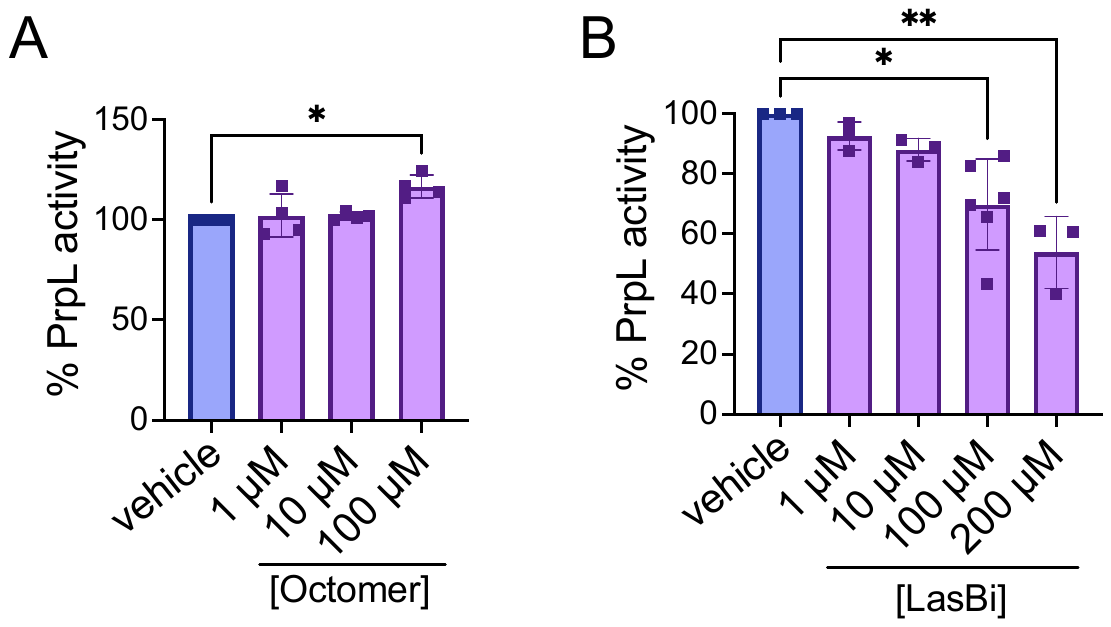

**Supplementary Figure 6. LasBi inhibits PrpL activity in a dose-dependent manner.** Percentage of PrpL activity relative to a vehicle control prepared with equivalent volume of DMSO. PrpL was pre-incubated for 15 minutes at 37 °C with increasing concentrations of A) octamer peptide or B) LasBi. PrpL activity was measured using Chromozym PL chromogenic substrate. Data are represented as mean ± standard deviation and were compared by one-way ANOVA with correction for multiple comparisons. Non-significant = p >0.05, * = p <0.05, and ** = p <0.01. Comparisons not otherwise indicated are non-significant.

**Supplementary Table 1. Clinical scoring criteria.**

| **Category** | **Score** | | | |
| --- | --- | --- | --- | --- |
|  | **0** | **1** | **2** | **3** |
| Physical appearance | No change | Mild ruffled coat and pulling back of ears | Moderate ruffled coat, occasional grimace | Very rough coat and grimace |
| Posture | Normal | Slight hunch | Moderate hunch | Severe hunch |
| Activity/behavior | Normal activity and response to stimulus | Slowed activity | Minimal activity | Responding only when stimulated |
| Respiration rate | Normal | Slightly elevated | Visibly elevated (rapid or laboured) | Very laboured (very rapid or very slow) |
| Body temperature (surface) | 29-34 °C | 27-28.9 °C | 25-26.9 °C | ≤ 24.9 °C |

A combined score of 12+ was an immediate humane endpoint for euthanasia, as were unconsciousness, inability to remain upright, agonal respiration, convulsions, and weight loss of over 15% of starting body weight.

**Supplementary Table 2. Antibodies used for western blotting analysis.**

| **Target** | **Catalog #** |
| --- | --- |
| c-Jun | 9165 |
| phospho-c-Jun | 9164 |
| c-Fos | 4384 |
| phospho-c-Fos | 5348 |
| IκB | 9242 |
| phospho-IκB | 2859 |
| GAPDH (HRP-conjugated | 3683 |

All antibodies were raised in rabbit and purchased from Cell Signaling Technology.

**Supplementary Table 3. Antibodies used for flow cytometry.**

| **Target** | **Fluorophore** | **Clone** | **Supplier** |
| --- | --- | --- | --- |
| MHCII | Brilliant Violet 510 | M5/114.15.2 | BioLegend |
| SiglecF | PerCP-eFluor 710 | 1RNM44N | Invitrogen |
| CD11c | Brilliant Violet 605 | N418 | BioLegend |
| CD11b | eFluor 450 | M1/70 | Invitrogen |
| CD103 | PE/Dazzle | 2E7 | BioLegend |
| Ly-6C | PE | HK1.4 | BioLegend |
| Ly-6G | PE/Cyanine7 | 1A8 | BioLegend |
| F4/80 | Alexa Fluor 700 | BM8 | BioLegend |
| CD45 | APC | 30-F11 | BioLegend |
| CD16/CD32 | N/A | S17011E | Biolegend |
